# High resolution ensemble description of metamorphic and intrinsically disordered proteins using an efficient hybrid parallel tempering scheme

**DOI:** 10.1101/2020.06.02.124628

**Authors:** Rajeswari Appadurai, Jayashree Nagesh, Anand Srivastava

## Abstract

Determining the conformational ensemble for proteins with multi-funneled complex free-energy landscapes is often not possible with classical structure-biology methods that produce time and ensemble averaged data. With vastly improved force fields and advances in rare-event sampling methods, molecular dynamics (MD) simulations offer a complementary approach towards determining the collection of 3-dimensional structures that proteins can adopt. However, in general, MD simulations need to either impose restraints or reweigh the generated data to match experiments. The limitations extend beyond systems with high free-energy barriers as is the case with metamorphic proteins such as RFA-H. The predicted structures in even weakly-funneled intrinsically disordered proteins (IDPs) such as Histatin-5 (His-5) are too compact relative to experiments. Here, we employ a new computationally-efficient parallel-tempering based advanced-sampling method applicable across proteins with extremely diverse free-energy landscapes. And we show that the calculated ensemble averages match reasonably well with the NMR, SAXS and other biophysical experiments without the need to reweigh. We benchmark our method against standard model systems such as alanine di-peptide, TRP-cage and β-hairpin and demonstrate significant enhancement in the sampling efficiency. The method successfully scales to large metamorphic proteins such as RFA-H and to highly disordered IDPs such as His-5 and produces experimentally-consistent ensemble. By allowing accurate sampling across diverse landscapes, the method enables for ensemble conformational sampling of deep multi-funneled metamorphic proteins as well as highly flexible IDPs with shallow multi-funneled free-energy landscape.

**Significance/Authors’ Summary:** Generating high-resolution ensemble of intrinsically disordered proteins, particularly the highly flexible ones with high-charge and low-hydrophobicity and with shallow multi-funneled free-energy landscape, is a daunting task and often not possible since information from biophysical experiments provide time and ensemble average data at low resolutions. At the other end of the spectrum are the metamorphic proteins with multiple deep funnels and elucidating the structures of the transition intermediates between the fold topologies is a non-trivial exercise. In this work, we propose a new parallel-tempering based advanced-sampling method where the Hamiltonian is designed to allow faster decay of water orientation dynamics, which in turn facilitates accurate and efficient sampling across a wide variety of free-energy landscapes.

## 1. Introduction

Biomolecules are not static but exhibit time-dependent dynamic motions that are tightly coupled with their functions. Molecular simulations are now increasingly relied upon for visualizing detailed events of such molecular motions at atomic resolutions, which are often not possible using experimental methods. Originally developed in 1950s,^1^ molecular dynamics (MD) simulation technique has come a long way since its first application on biological molecule in 1977.^1,2^ With wide-range of applications, MD simulations are now routinely employed in important biomolecular characterization such as stability, flexibility, conformational changes, ligand binding and the effect of perturbations upon mutations, post-translational modifications and environmental changes.^3,4^ Increased availability of high-performance computers in the last decade further expands its accessibility and allows simulating processes that take longer than microsecond timescales^5,6^ and involve billions of atoms.^7,8^ Nonetheless, sampling in classical MD simulations is limited to the local minima of the free energy landscape. Consequently, accessing the complex energy landscapes of proteins is far from trivial for proteins with multi-step folding,^9,10^ metamorphic proteins,^11–13^ and intrinsically disordered proteins (IDPs).^14–16^

Enhanced molecular dynamics simulations such as temperature replica exchange molecular dynamics (TREM),^17^ are particularly useful in sampling conformations across energy barriers and therefore expediting the observations of rare biomolecular transitions. In replica exchange method, conformations are sampled using multiple replicas simulated at a series of low and high temperatures and are stochastically exchanged at regular intervals to provide an unbiased Boltzmann-weighted ensemble of conformations. However, the number of replicas required for the simulation scales exponentially with the degrees of freedom of the system hampering its application in large biomolecular solutions. Bruce Berne and coworkers developed a powerful alternative called replica exchange with solute tempering (REST)^18,19^, where they designed the Hamiltonian to partially heat a subset of the system (solutes). The approach reduced the required number of replicas by multiple folds. Such a powerful design of the Hamiltonian has been successfully shown to simulate weak binding of Aβ peptide on a lipid bilayer,^20^ lateral equilibration of lipids in a bilayer^21^, and a 90-residue long SH4 unique domain protein.^22^ While a much wider applications in protein folding is envisioned, however, the method underperforms in proteins where the conformations are separated by large free energy barriers producing poor mixing of replicas between the high temperature regime and low temperature regime. This is likely due to the discrepancy between the energy compensation of hot solute and cold solvent. Inclusion of a subset of random water molecules to the tempering central group improves the mixing but at the expense of poor scaling with the system size.^23^ More recently, a generalized version of the REST (gREST) that scales the Hamiltonian based on particles as well as energy terms has also been introduced,^24^ which further reduces the number of replicas and alleviates the scalability issues to a large extent. However, as the extent of tempering is limited to few energy terms, the method requires longer time to converge on the conformational sampling. This suggests that crossing of the free energy barrier is still a bottle neck for this class of methods.

In general, parallel tempering methods suffer when the barriers between folded/unfolded states and intermediate states are high or when the transition state is diffusive in nature with the large entropic barrier slowing down conformational exchanges.^25–27^ The unrealistically reduced residence time in the metastable states prevents switches between conformational basins. In these cases, friction between the polypeptide chain and solvent modulates the speed of protein folding or unfolding.^28^ In this work, we propose a hybrid replica exchange method (termed hereafter as Replica exchange with hybrid tempering, REHT) that differentially and optimally heats up both the solute and solvent. In other words, along with the Hamiltonian scaling of each replica that effectively heats up the protein solute, the replicas are also coupled to different high temperature baths that heat up the system including solvent particles. The chosen bath temperature range of the replicas is small enough such that the energy difference due to solvent self-interaction is minimal and does not lead to the scalability issue. At the same time, the optimal tempering of the solvent along with the solute ensures an efficient rewiring of the hydration shell that work in cohort with that of protein conformational change, thereby helping in crossing larger barriers and in particular the entropic barriers. We demonstrate this by applying the protocol on a diverse set of proteins that differ widely on their size and complexity of the underlying free energy landscape. The choice of protein molecules ranges from simple model system such as alanine dipeptide to fast folders such as TRP-Cage and β-hairpin and proteins with complex energy landscapes such as IDPs (example: Histatin-5) and metamorphic proteins (example: RFA-H).

## 2. Results and discussion

We explore the multidimensional free energy landscapes of diversely complex proteins using the newly designed REHT method and compare its efficiency with that of the state-of-the-art REST2^19^ simulations. Towards this, we exploited the HREX module of PLUMED, originally developed for performing the Hamiltonian replica exchange simulations.^29^ The module is very flexible and allows for simultaneous use of different bias in the replicas such as the Hamiltonian, collective variable, temperature and pressure. For REHT method, we include the additional temperature bias in the replicas along with the Hamiltonian scaling of the protein solute and for which we derive and apply the corresponding detailed balance exchange criteria. The derivation of the exchange criteria, honoring detailed balance, is provided in the supporting information (SI). In short, the exchange criteria for REHT is given by:

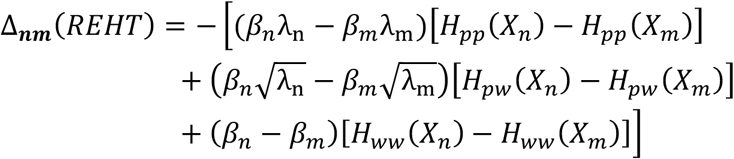

where, *H*_*pp*_(*X*_*m*_) and *H*_*pp*_(*X*_*n*_) indicates the intramolecular energy of protein solute in m^th^ and n^th^ replicas, respectively. *H*_*pw*_(*X*_*m*_) and *H*_*pw*_(*X*_*n*_) are interaction energies between protein and water sites in the two replicas. Similarly, *H*_*ww*_(*X*_*m*_) and *H*_*ww*_(*X*_*n*_) indicates the water-water interactions in the corresponding replicas. *β*_*m*_ and *β*_*n*_ are inverse temperatures and λ_m_, λ_n_ are Hamiltonian scaling of replica m and n, respectively.

Though the methodological advancement of REHT has its origin in REST2, the new method has magnificent impact on the sampling efficiency as will be demonstrated here in this section. The efficiency is not just shown on simple model proteins but also on proteins such as intrinsically disordered Histatin-5 and metamorphic RFA-H proteins. The different systems considered in this paper are shown in Fig. 1. The computational details of all the systems are presented in Table S1.

**Figure 1:**
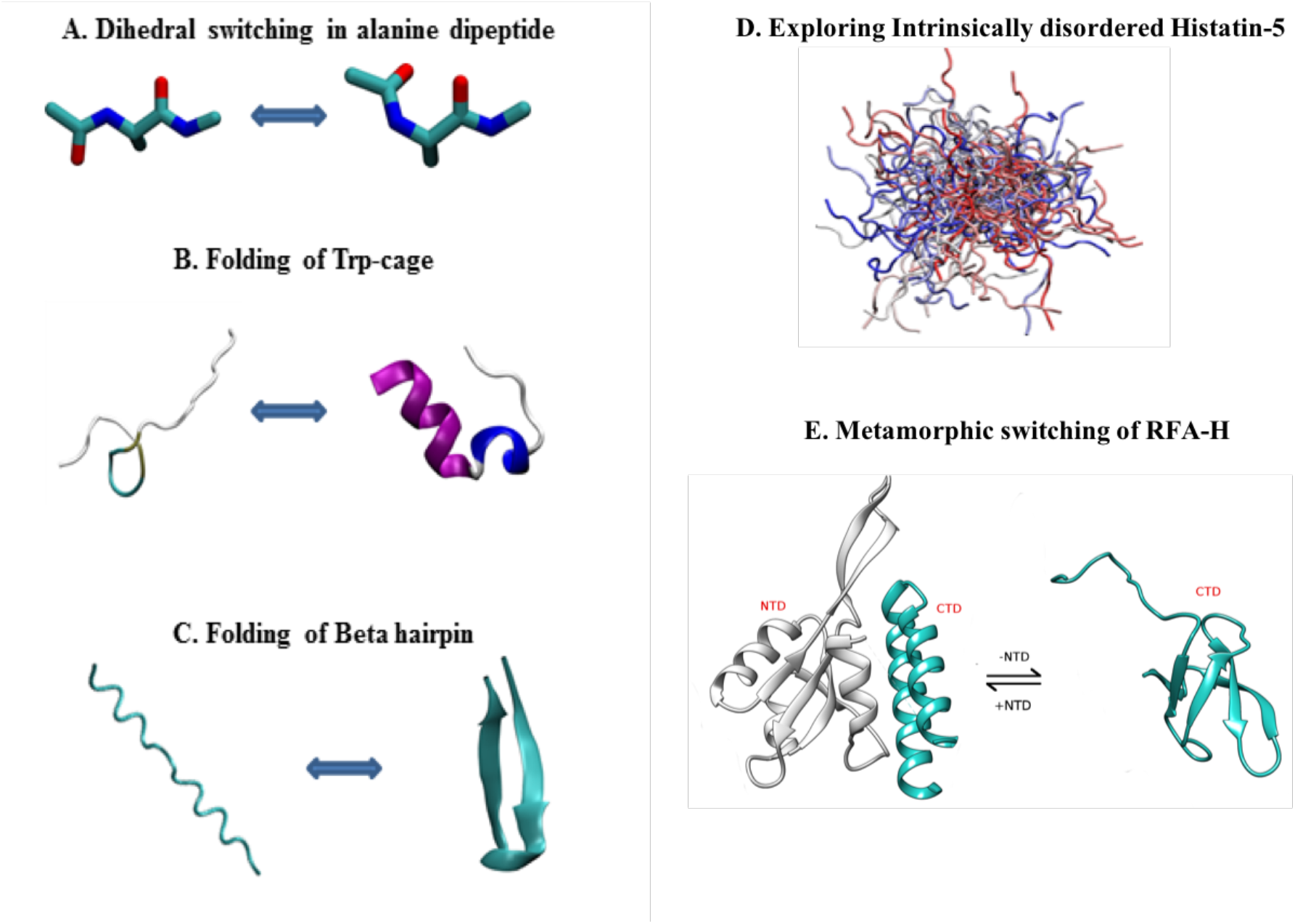
List of systems studied: A) Dihedral switching of alanine dipeptide, B) Folding of Trp-Cage from completely unfolded structure, C) Folding of β-hairpin, D) Intrinsically disordered Histatin-5, and E) Metamorphic switching in bacterial RFA-H are explored using the state-of-the art REST2 and the newly designed REHT approach.

### 2.1 Modulated tempering of water self-interaction promotes conformational search

We first test the sampling efficiency of our parallel tempering method on well-known model systems alanine-dipeptide, TRP-cage and β-hairpin. Though TRP-cage and β-hairpin are widely recognized with two-state folding kinetics,^30^ several experimental and simulation studies support the presence of intermediate states indicating the complex nature of folding energy landscapes of these fast-folders.^31,32^ For our simulations, the completely extended starting conformations of physiologically relevant zwitterionic state of TRP-cage^33^ (with charged termini) and β-hairpin were obtained by simulating them at 600 K in explicit water for 10 ns timescale. For comparison against the state-of-the-art in MD-based sampling, the simulations are also performed with REST2.^18,19,22^ Replica mixing of two simulations in both the systems are shown in Fig. S1 and S2.

During the course of the simulations, both the REHT and REST2 methods drive the transition of the unfolded starting conformation to a native folded conformation with almost a perfect match with the experimentally found native structure (0.4 Å RMSD) as shown in Fig. 2A and Fig. S3A. However, we observe faster transition with REHT between the folded and unfolded basins. The time evolution of RMSD indicates REHT method samples the folded structures of the two proteins in less than 100 ns timescales (Fig. 2B) and produces the folded state in 5 out of 12 replicas (Fig. S3B, S4). The REST2 simulations sample the folded structure of TRP-cage (Fig. 2B) and β-hairpin (Ref^24^) at around 300 ns. Moreover, only 1-2 replicas out of 8 replicas are independently folded in these simulations (Fig. S5 and Ref^24^). Free energy landscapes, as functions RMSD and radius of gyration, of the conformations sampled by the lowest temperature replica of TRP-cage simulations are presented in for REHT and REST2 in Fig. 2C and Fig. 2D, respectively. The predicted free energy barrier by the REHT method (~2 Kcal/mol) matches closely with the suggested free energy barrier of ~ 2.1 Kcal/mol,^30,34^ while a larger barrier of about ~ 6 Kcal/mol is observed in REST2 simulations. Further, the time evolutions of the one-dimensional free energy landscape along a single reaction coordinate, RMSD, (Fig S6) reveals that the REHT converges much faster than the REST2.

**Figure 2:**
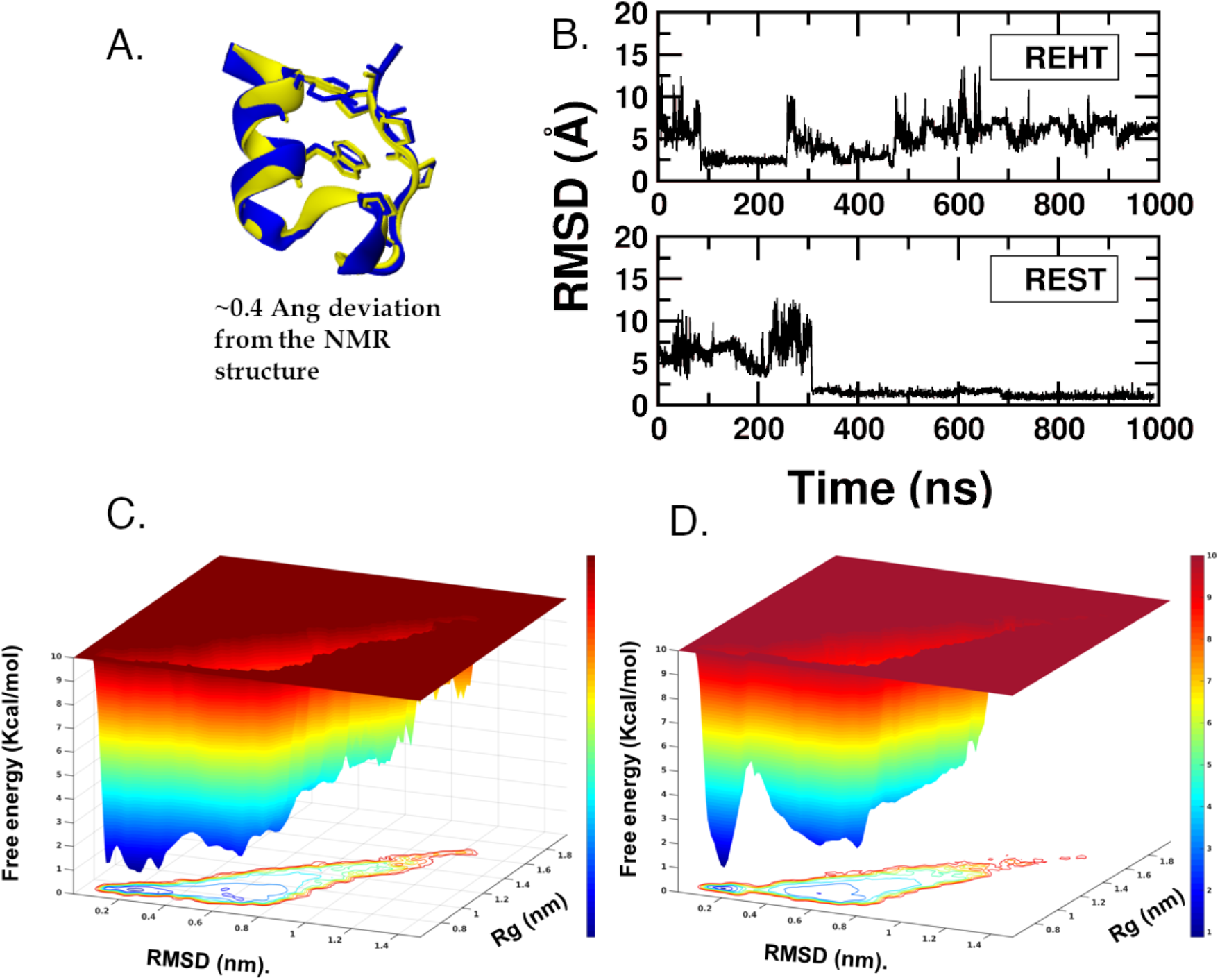
Efficiency of REHT in comparison to REST2 in capturing the folding of fast-folding protein, Trp-Cage. (A) Structural overlay between the natively folded NMR structure (blue) and the REHT-generated folded structure of Trp-cage (yellow, obtained at the base replica) (B) Time evolution of protein backbone RMSD from the NMR structure along one of the successfully folded replicas of REHT (top) and REST2 (bottom) simulations. The RMSD evolution of other folded replicas are shown in Fig.S3 and S4. (C) and (D) Free energy landscape of Trp-Cage, shown as a function of Radius of gyration (Rg) and RMSD against the NMR structure. The landscape is shown for the ensembles collected at the base replica of C) REHT simulation and D) REST2 simulation.

The origin of the difference lies in the way water self-interaction is treated in REHT and REST2. In Fig. 3, we present the water reorientation dynamics and H-bond dynamics in REST2 and REHT simulations. Fig. 3A clearly shows that the water reorientation is much faster in REHT method than the REST2 method. This difference is observed in almost all hydration shells and particularly evident in the first hydration shell. By optimally heating up the water, our method improves the orientation dynamics of water at the hydration layer. Consequently, this leads to the rewiring of H-bonds between protein and water measured here as an autocorrelation function of H-bond lifetime (Fig 3B). As illustrated, the H-bonds decay much faster in the new method in comparison to the REST2 method. The faster decay in the water orientation and H-bond allows for efficient rewiring of proteins allowing for the faster and more faithful conformational transitions.^35^

**Figure 3:**
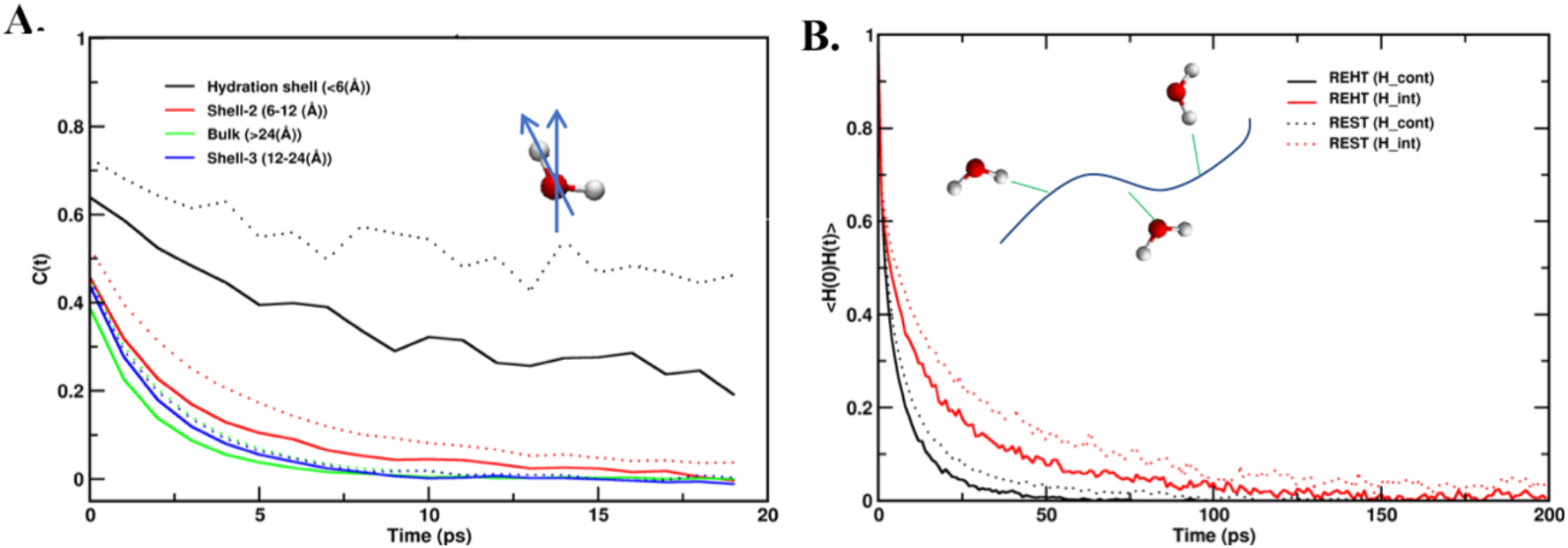
Extent of hydration dynamics and the resulting changes in the lifetime of protein-water H-bonds in Trp-cage folding: A) Average orientational autocorrelation function of water molecules (for OH bond vector) at different hydration shells surrounding the surface of protein. B) Time correlation functions of protein-water H-bonds formed by both continuous (black) and intermittent (red) water molecules. In both A and B, solid line represents the REHT simulation and dotted line represents the REST2 simulation.

This expedited sampling of the “slow” reaction coordinate (collective variable) is also visible in alanine dipeptide, the quintessential model system for barrier crossing events in biophysical advanced sampling methods development field. We discuss this in detail in SI under the subheading *“Dihedral switch in Alanine dipeptide”.* In short, we show that relatively slow transitioning ϕ-angle of the Ramachandran map^36,37^ is frequently sampled in the REHT method (Fig. S7-S8) with a larger number of replicas (4 out of 5) exhibiting this transition as well.

### 2.2 Ensemble sampling of flexible IDPs with high charge and low hydrophobicity

Accurate conformational sampling of IDPs is a major challenge in the field of molecular simulations where conventional forcefields do not seem to accurately reproduce the properties of IDPs. Several solutions towards fixing the forcefield transferability issues have been suggested and applied extensively to IDPs.^38–43^ What stands out in these improvements is the necessity to have a balanced protein-water interactions besides the required changes in the parameters of proteins.^44–46^Also, rare-event sampling methods are frequently used in conjunction with these improved force-fields to faithfully capture the conformational landscape of IDPs as shown for P53,^47^ alpha synuclein,^48^ islet amyloid polypeptide,^49^ amyloid beta,^50^ and NCBD IDPs.^51^ Recently, REST2 in combination with the IDP-specific forcefield (Amberff03ws) and water model (TIP4P/2005s) was used to successfully yield experimentally consistent ensemble in SH4UD.^22^

Despite these successes, there are set of IDPs on which the simulations data show striking deviations from the experimental results. For instance, in a 24-residue long antimicrobial peptide Histatin-5 (His-5), the simulation-generated ensemble deviates substantially from the experimental data from CD, NMR and SAXS measurements.^52,53^ His-5 fails to be accurately sampled even with the state-of-the-art forcefields^52,53^ suggesting the lack of transferability across IDPs with diverse range of charge and hydropathy characteristics. We illustrate the charge-hydropathy plot of various IDPs in Fig 4A. The plot clearly shows that the successfully characterized proteins (p53, AB42, NCBD, SH4UD) fall at or near the line separating the folded and unfolded region indicating the pre-molten globular nature of these IDPs. His-5 on the other end is located at the far end of charge-hydropathy plot with very minimal hydrophobicity and higher charges.

**Figure 4:**
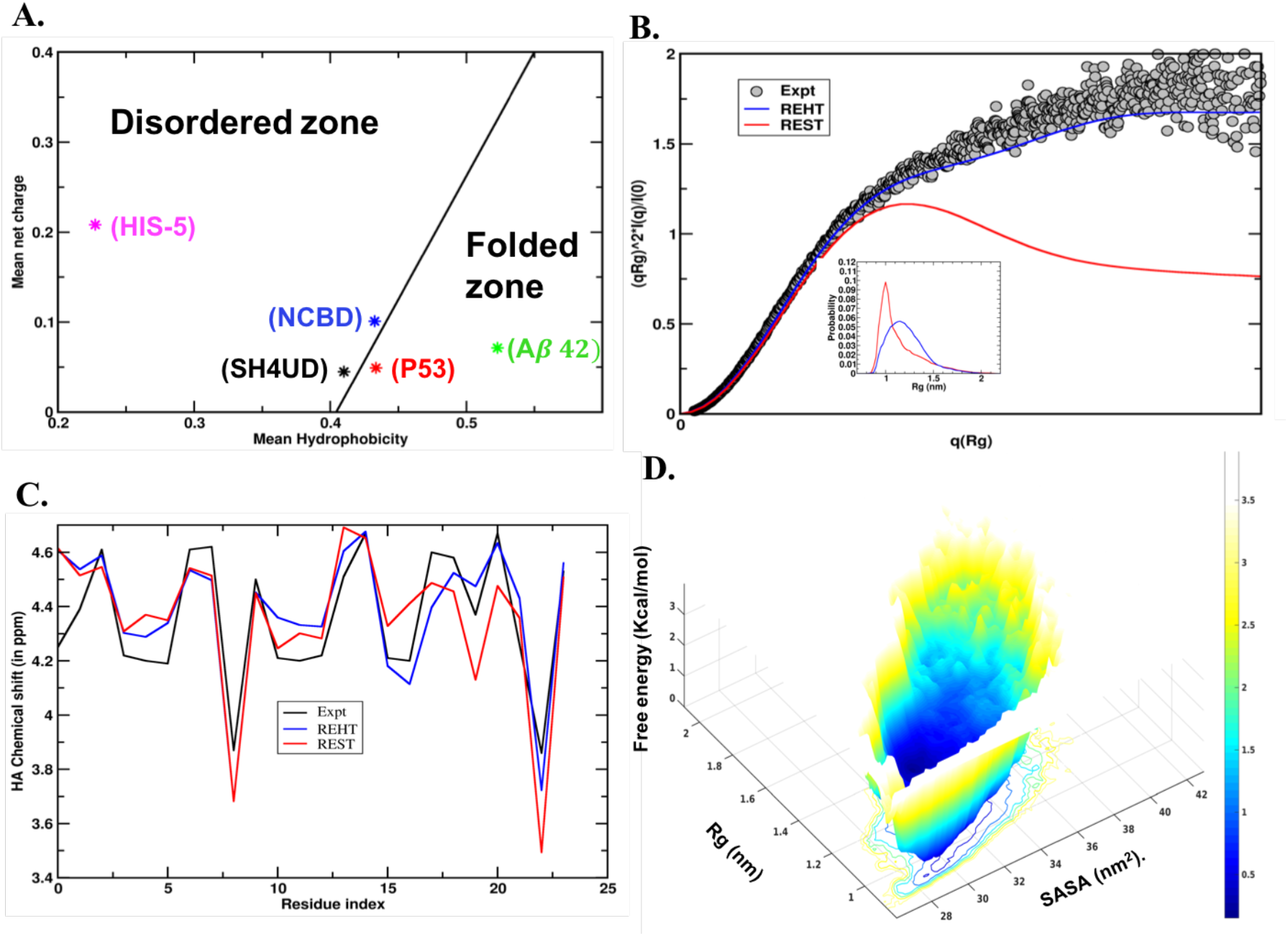
Ensemble description of Intrinsically disordered Histatin-5: A) Charge hydropathy plot showing the uniqueness of Histatin-5 that locate at the extreme disordered zone unlike other successfully-studied IDRs which exist at or near the folded zone. B) Comparison between the experimental (black) and theoretical ensemble-averaged SAXS profiles, represented as Kratky plot. The theoretical prediction was made for the last 250ns unweighted trajectories corresponding to the base replica of REHT and REST2 simulations. The distribution of Rg for the ensembles obtained from REST2 (red) and REHT (blue) simulations are shown in the inset. C) Comparison of ensemble averaged chemical shifts of H-alpha atoms predicted from the REHT and REST2 simulations (for the same 250ns trajectory) with reference to the experimental NMR chemical shifts. D) Weakly funneled diffusive energy landscape of Histatin-5 shown as a function of Rg and solvent accessible surface area.

We sampled the conformational landscape of His-5 using REHT and REST2 (see Fig. S9 for mixing of replicas) and verified the accuracy of the generated ensemble against various experimental ensemble average properties. To start with, we evaluated the secondary structure content of this peptide by analyzing the backbone dihedrals of all the residues. Circular dichroism data^54^ reveals that His-5 prefers to form polyproline II (PPII) structures and the PPII propensity is lost at higher temperatures. Fig. S10 in SI shows the Ramachandran map for His-5 indicating the most probable occurrence of PPII structures, which is generally not recapitulated in the conventional IDP sampling methods.^54^ Interestingly, enhancing the sampling either by REHT or REST2 recapitulates the propensity of PPII structures of His-5 in a qualitative agreement with the CD measurements. Furthermore, the temperature dependent loss of PPII structure is also captured correctly (Fig S10).

We also calculate the nanoscale structural properties of His-5, often measured experimentally by means of small angle X-ray scattering (SAXS) methods that describes the overall size and shape of a protein in solution. The experimental SAXS data^55^ for His-5 at room temperature and at neutral pH was obtained from Prof. Marie Skepo’s laboratory in Sweden. The results are depicted in Fig. 4A as dimensionless Kratky representation, that qualitatively assess the compactness and flexibility of protein. This can be obtained from the form factor using the following equation: (*qRg*)^2^*I* (*q*)/*I* (0) and plotted against *qRg*. In case of well folded proteins, the Kratky plot exhibits a bell-shaped peak at low-q regime and converges to the q-axis at high-q regime, whereas in IDPs it shows monotonic increase in intensity at the high-q region.^56,57^ As shown in the figure, the Kratky plot of His-5 obtained from the experiment exhibits the signature of highly flexible and extended IDP. The SAXS profile from REHT ensemble data (Fig. 4B) matches closely with that of the experiment thereby reinforcing its fidelity in constructing the IDP ensemble accurately. We believe that conformations of IDPs that lie significantly away from the CH border area are far more affected by the hydration-induced stability. In such cases, REST2 is not suitable since it does not treat the surrounding water and hence produces a more compact protein configurational ensemble as evident from both the SAXS profile and Rg plot (Fig 4B:inset). We also compare the chemical shifts of hydrogen atoms linked to the Cα predicted from the REHT and REST2 methods with that of the NMR chemical shifts data (Fig. 4C). The chemical shifts predicted from REHT matches better with the experiments than the REST2 method.

After verifying the REHT-generated ensemble with the experimental data from CD, SAXS and NMR measurements, we reconstitute the free energy map of His-5 at the room temperature as a function of various structural parameters (Fig. 4D, Fig S11-S12). The free energy map indicates rather a flat-bottom low-energy landscape across a wide range of Rg (from 1.0 nm to 1.8 nm) and SASA values (28 nm^2^ to 38 nm^2^). The results indicate that under solution conditions, the peptide prefers to exist in a completely disordered conformation. Such a large disorderliness endows them with an ability to bind diverse targets as confirmed from the proteomic analysis.^58^ We also traced for the residual alpha helical propensity as observed when the peptide approaches the biological membrane during its candidacidal functioning.^59^ However, we did not observe any trace of helicity in the solution ensemble (Fig. S13), suggesting the helical transition may be adapted in an induced fit manner upon association with the membrane.

### 2.3 Deciphering the transition intermediates of metamorphic protein (RFA-H)

Unlike the weakly-multi-funneled landscapes generally found in IDPs, metamorphic proteins have deep multi-funnels with each low energy basin representing a distinct well-folded conformation. These conformations switch upon being signaled to perform different functions. Among the many members of metamorphic proteins known till date, RFA-H possesses the highly divergent folds with completely different secondary structures. RFA-H is a bacterial regulatory protein, in which its C-terminal domain alternates between all α-helical and all β-sheet conformations based on presence or absence of inter-domain contacts respectively (Fig 5A).^60^ While in helical conformation, it activates the transcriptional elements, in beta sheet conformation it recruits ribosomal protein and thereby couples to translation.^61,62^ Transition between these alternate conformations generally involve unfolding from one conformation and refolding to other conformation.^63^ Exploring these protein landscapes by the computational MD simulations is highly challenging and rarely attempted utilizing biasing techniques such as targeted MD or assisted GO model.^64–66^

**Figure 5:**
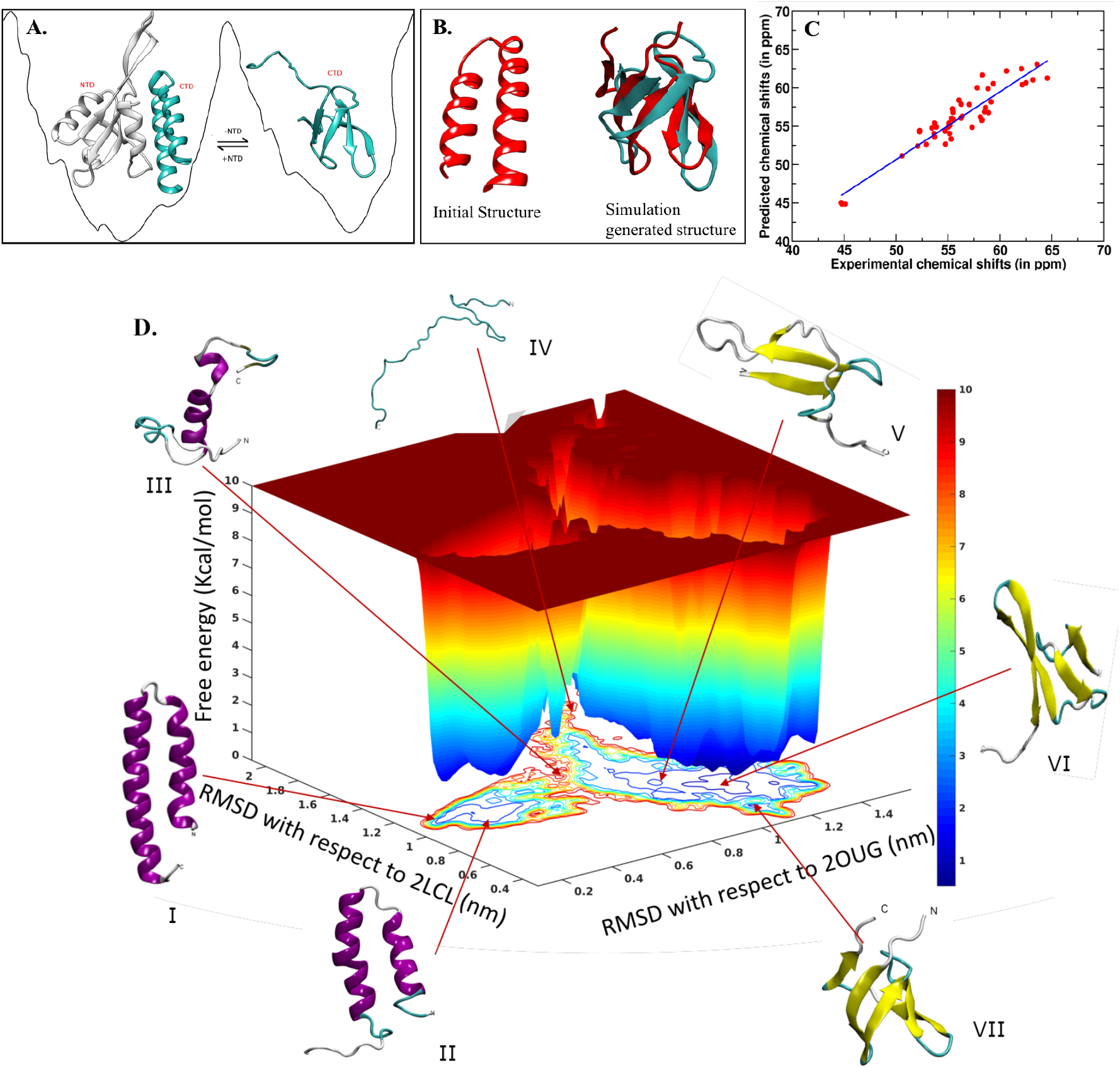
Conformational metamorphosis of RFA-H: A) Schematic representation of the two native structures resolved in experiments, the α-helical fold of CTD in presence of NTD and beta fold in isolated CTD. B) The initial and final structures of the REHT simulation. The helical state of CTD while deleting the NTD was chosen as the starting structure (left figure in B). Structural superposition of the final simulation generated beta barrel structure (red) over the experimental beta structure (cyan) is shown on the right side of (B). C) Validation of simulation generated ensemble by comparing the predicted C-alpha chemical shifts with the experimental shifts. The fitted linear regression (blue line) indicates a precise match between the two sets of data. D) Dual basin free energy landscape of RFA-H shown as a function of RMSDs from the experimentally found α-helix (2OUG) and beta-sheet (2LCL). The conformations at each of the basins, all α-helical and all β-sheet (I and VII), early structure with the loss of helicity at both the termini (II), intermediate partially unfolded structure with residual α-helical content (III), and metastable structures with open beta sheets (V and VI) are shown along the landscape. Note the completely unfolded structure (IV) is at the other side of the dual basin and the transition of α → β structure doesn’t need to step through the completely unfolded structure unlike that of metamorphic lymphotactin.^63^

We set to explore the conformational metamorphosis of RFA-H in explicit water using our unbiased REHT simulation. The same was attempted using REST2 simulation as well, which did not yield proper mixing of replicas even at 250 ns of simulation time per replica (Fig S14) and hence was dropped. REHT started showing mixing between replicas by around 100 ns and the runs were extended to 1000 ns/replica for the 25 replicas. Experimentally obtained α-helical state (PDB ID: 2OUG) of the C-terminal domain (residue number 115-162) was used as starting structure (Fig. 5B). Unlike the previous studies using targeted MD^64,66^ or assisted GO model with replica exchange tunneling,^65^ the REHT method doesn’t embed in itself any information of the target state. In spite of that, the simulations spontaneously transform the RFA-H_CTD_ from helical structure to the beta barrel conformation. Secondary structure analysis of the ensemble generated at the lowest temperature (310K) replica reveal that this transformation occurs within 300 ns timescale (Fig. S15). The residues that were shown to form beta sheets agree well with the NMR-generated structure (PDB ID: 2LCL). The structural overlay of a snapshot obtained from the simulation on to the NMR structure is shown in Fig. 5B and their root mean squared deviation is calculated to be 3.4 Å. It should be appropriate to note here that the previous unbiased MD simulation, with implicit solvent, fails to produce the beta barrel spontaneously. Additional refinement with explicit solvent simulation was required to arrive at the barrel structure with ~5 Å RMSD from the experimental structure.^67^

To further validate our results, we predicted the backbone chemical shifts of the simulation-generated ensemble obtained from the room temperature replica. The last 500 ns timeframes without any reweighting or constricted selection was used for this analysis. The predicted C_α_ and C_β_ chemical shifts were compared with those measured from NMR spectroscopy (Fig 5C). The figure clearly demonstrates a striking agreement between the predicted and measured chemical shift values. Following the experimental validation, we traced for the molecular details of the fold switching mechanism from our atomic-resolution trajectory, the information that cannot be obtained directly from the experiments. Towards this, we plotted the free energy map as a function of RMSDs with respect to the experimentally found helical and beta-barrel structures (Fig 5D). The map reveals dual-basins, each corresponding to the different folds of the RFA-H, where the basins are separated by a free energy barrier of about 5 Kcal/mol. This barrier corresponds to the α→β transition while not favoring the reverse transition. This argument is based on our observation with increasing simulation time, the population of β conformation shows an increase suggesting a (non-converged) deeper basin for the β structure. (Fig S16) For the reverse transition β→α, experiments indicate that the N-terminal interaction is important.^60^ To reinforce this statement we simulated additional simulation initiated from the NMR-obtained beta structure, which show stable beta barrel structures (Fig. S17), and is consistent with the NMR observations.^60^

While the free energy basin corresponding to the α-helical structure exhibits a funnel like architecture, that of the β-barrel structure looks like a rugged landscape indicating possible heterogeneity in that state. The heterogeneity likely builds up the entropic barrier and adds to the complexity of the free energy map and thereby limits the spontaneous transition from one state to the other upon using conventional advanced sampling methods. REHT shows promising results in crossing both the enthalpic and entropic barriers and therefore allows for studying the complex conformational transitions of IDPs and “transformer” proteins such as RFA-H. The method also opens up avenues to study other transformer proteins such as chemokine lymphotactin protein,^68^ Mad2 spindle checkpoint protein,^69^ CLIC1.^70^

## 3. Materials and Methods

### 3.1 Parallel tempering

For the sake of completion, we provide the theoretical background about the different versions of replica exchange methods in SI under the subheading “*Replica exchange simulation methodology*”. Moreover, the detailed balance exchange condition for the newly developed REHT method, with a specially designed Hamiltonian that differentially treats solvent and solute across a temperature range, is also derived and presented in SI.

### 3.2 Systems and simulation details

#### 3.2.1 List of systems studied and preparation of atomistic model

Fig. 1 and Table S1 contain information about the diverse sets of proteins used in this work. The initial unfolded structure of Trp-Cage and β-hairpin (C terminal hairpin of B1 domain in protein G) was obtained by simulating the NMR structures (PDB Id: 1l2Y and 1lE3, respectively) at 600K temperature for 10 ns. For Hist-5, the unfolded structure was built using VMD protein builder by feeding in the sequence information. Simulation of RFA-H C-terminal domain was initiated from the α-helical conformation as obtained from the X-ray crystallography (PDB ID:2OUG) used as initial structure for the simulation. All the proteins (system 1-5 in Fig.1) were solvated in a cubic box with a minimum distance of 1.2 nm from the surface of the protein (in case of RFA-H 1.5 nm is used). 3-site rigid TIP3P water model was used commonly for all the systems. The systems were also neutralized with physiological concentration of NaCl (0.15M). For the topological parameters of fast folding proteins, we used Amberff14SB in order to faithfully compare the efficiency with earlier REST2 and gREST simulations,^24^ whereas for all the rest of the systems (alanine dipeptide, His-5 and RFA-H), Charmm36m^39^ force field was used.

#### 3.2.2 Details of MD simulation

The modelled proteins solutions were initially energy minimized using steepest descent algorithm for 50,000 steps to avoid any poor contacts. The minimized structure was then thermalized and equilibrated sequentially in NVT and NPT ensembles each for 2 ns. The protein and the solvent were coupled separately to the target temperatures using modified Berendsen thermostat. The pressure was coupled at 1 bar using Parrinello-Rahman barostat. The final production simulation was performed in NVT ensemble using Nose-Hoover thermostat. A cut-off of 1 nm was used for calculating the electrostatics and VdW interactions. Particle Mesh Ewald was used for long-range electrostatics. Leap-frog integrator with the time step of 2 fs was used to integrate the equations of motions. All the hydrogen atoms were constrained using LINCS algorithm. The simulations were performed using Gromacs-2018.2 patched with Plumed-2.4.1. All the parameter files were uploaded in the GitHub repository (https://github.com/codesrivastavalab/ReplicaExchangeWithHybridTempering)

#### 3.2.3 Replica exchange parameters

For all simulations, we ran both our newly designed REHT scheme as well as the state-of-the-art REST2 method. The two methods differ in the way the protein and water molecules are treated in the Hamiltonian. REST2 tempers only the protein but keeps the water at room temperature by scaling the potential energy function of solute. The bath temperature used in REST2 is constant across all the replicas. Whereas in REHT method the bath temperature is raised mildly up to 340K as we go up on the replica ladder. This sufficiently heats up the water molecules for its efficient dynamics in REHT method. For enhancing the protein dynamics, the potentials of the protein solute are scaled up to a maximum factor (λ) of 0.5 similar to the REST2 method. To ensure the faithful comparison, the overall effective temperature realized on the protein is identical in both the simulations (Table. 1). However, due to the extra degrees of freedom due to treatment of the water, the number of replicas used is moderately higher for the REHT method than for the REST simulations. More details and the scripts for all the input preparation (including the scaling of Amber14SB and charmm36m forcefields) are given in the github link. (https://github.com/codesrivastavalab/ReplicaExchangeWithHybridTempering)

### 3.3 Conformational ensemble analysis details

#### 3.3.1 Theoretical SAXS analysis

Theoretical SAXS profile for the HIS-5 ensemble was predicted using CRYSOL^71^ that calculates orientationally averaged scattering pattern using multipole expansion from atomic coordinates while considering the solvation shell using spherical harmonics. The predicted value is compared against the experimental form factor of His-5 obtained at neutral pH from Skepo et.al.^55^

#### 3.3.2 Theoretical analysis of chemical shifts

The backbone chemical shifts of RFA-H atomic coordinates were calculated using SPARTA+,^72^ that works based on artificial neural network. To validate our simulation, the predicted shifts were compared against the experimental NMR chemical shifts as obtained from BMRB database entry 17615.^60^

#### 3.3.3 Structural analysis

Other structural parameters such as root mean squared deviation (RMSD), radius of gyration (Rg) and backbone dihedral angles were analyzed using the respective Gromacs utilities. Analysis of protein secondary structures was performed using DSSP tool. VMD was used for visualizing the trajectory and rendering of protein cartoon images.

#### 3.3.4 Analysis of hydration dynamics

The dynamical properties of the hydration shell were calculated by means of water orientational relaxation and hydrogen bond lifetime decay as implemented in MDAnalysis package.^72^ Continuous trajectory with respect to simulation time was used for this analysis. The orientational relaxation essentially indicates the rotational freedom of water molecules in a solvation shell. This is measured by computing the second order rotational autocorrelation function of two vectors: one along the OH bond and the other along the dipole moment of water, as given as: 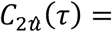 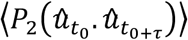, where, *P*_2_(*x*) is the second Legendre polynomial and 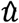 is the unit vector. Similarly, we measure the persistence of interaction between the protein and the water molecules by computing the autocorrelation function of Hydrogen bond lifetime, as follows: 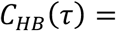 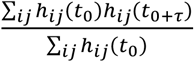, where, the H-bond between pair i and j at time t, (*h*_*ij*_(*t*) is considered to be 1 if the geometric distance (< 3.5 Å) and angle (>120°) cutoff are met. *h*_*ij*_(*t*) = 0 otherwise. We considered two types of protein-water H-bonds: ‘continuous H-bond’ when a water molecule is continuously involved in H-bonding and ‘intermittent H-bond’ when the H-bond is retained with intermittent change of water molecules.

#### 3.3.5 Calculations of Free energy change

The relative Gibbs free energy of an equilibrium ensemble is computed as a function of two reaction coordinates as follows:

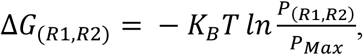

Where *K*_*B*_ represents the Boltzmann constant, T is the temperature. *P*_(*R*1,*R*2)_ denotes the probability along the two reaction coordinates, which is calculated using a k-nearest neighbor scheme and *P*_*Max*_ denotes the maximum probability. The 3-dimensional representation of the free energy surface was plotted using Matlab.

## 4. Conclusions

In this work, we design a new scaled Hamiltonian that differentially treats the protein and solvent interactions as a function of temperature in the replica-exchange framework. Essentially, the design of the Hamiltonian in our method (REHT) allows faster decay of water orientation dynamics, which in turn enhances the conformational sampling for proteins. We find that the accelerated thermodynamic sampling in REHT compensates for the additional computational cost incurred by the moderately higher number of replicas due to inclusion of water interactions in the redesigned Hamiltonian for replica exchanges. The high-resolution structural ensemble for a “variety of proteins” produced through our REHT simulations agree excellently with the ensemble average observables obtained from NMR, SAXS, CD and other biophysical experiments without the need of any reweighting. The method is particularly suited for highly flexible IDPs such as His-5 where solvation dynamics stabilize and drive the coexistence of multiple degenerate extended states of the peptide on a free-energy surface that have multiple shallow basins. Large free energy barriers in metamorphic proteins, with multiple well-defined basins having diversely folded conformations, are usually not easily surmountable through conventional methods. We show that REHT is capable of sampling across the barriers of metamorphic proteins, deduce possible transition intermediates and solve their conformational ensemble. As a future perspective, we believe that when appropriately coupled with the known experimental data, the high-fidelity structural ensemble information from the unconstrained REHT-simulations can be effectively used to sample and if needed further minimize the residual with experimental observables using integrative modeling framework.^73–75^

## 5. Acknowledgments

The authors are grateful to the SciNet HPC Consortium, ComputeCanada^76,77^ for the computational resources and Prof. Marie Skepo for the Histatin-5 SAXS data. A.S. and A.R. thanks Dr. Jagannath Mondal, Dr. Ashok Sekhar and Dr. Debostuti Ghoshdastidar for their valuable comments and suggestions on the manuscript. AR thanks Wellcome Trust – DBT India Alliance for Early Career fellowship. J.N. thanks DST-SERB for funding under Ramanujan Faculty Fellowship. A.S. also thanks IISc-Bangalore and the Ministry of Human Resource Development of India for the startup grant and the Department of Science and Technology of India for the early career grant. This research was also supported by the Department of Biotechnology, Government of India in the form of IISc-DBT partnership program. Support from FIST program sponsored by the Department of Science and Technology and UGC, Centre for Advanced Studies and Ministry of Human Resource Development, India is gratefully acknowledged by the authors.

## Supporting information

**Table S1:** System descriptions and details of the replica exchange simulation parameters.

| System studied | Number of atoms | Number of replicas | Overall Effective temperature range | $\lambda$ scaling and bath temperature | Cumulative time (in ns) | Average exchange acceptance probability | Transit time<sup>#</sup> (in ns) |
| --- | --- | --- | --- | --- | --- | --- | --- |
| Alanine dipeptide | 1864 | 5 (3) | 300–600K (300–600K) | $\lambda=1-0.567$; T=300-340K ($\lambda=1-0.5$; T@300K) | 100 (60) | 0.25 (0.30) | 6 (4) |
| Trp-cage | 21038 | 12 (8) | 300–478K (300–478K) | $\lambda=1-0.71$; T=300-340K ($\lambda=1-0.63$; T@300K) | 12000 (8000) | 0.12 (0.30) | 10 (2) |
| Beta-hairpin | 17203 | 12 (8*) | 300–478K (300–478K) | $\lambda=1-0.71$; T=300-340K ($\lambda=1-0.63$; T@300K) | 12000 | 0.12 | 5 |
| Histatin-5 | 23195 | 15 (10) | 300–478K (300–478K) | $\lambda=1-0.71$; T=300-340K ($\lambda=1-0.63$; T@300K) | 11250 (7500) | 0.15 (0.45) | 3 (1) |
| RFA-H | 40861 | 25 (25) | 310-705K (310-705K) | $\lambda=1-0.48$; T=310-340K ($\lambda=1-0.44$; T@310K) | 25000 (6250) | 0.17 (0.45) | 110 (Not achieved till 250ns) |

Primary values correspond to that of REHT simulations, and the values in bracket is for REST2 simulation.
\* In case of Beta-hairpin the REHT simulation is compared with the REST2 simulation performed earlier (Ref:1) using the same forcefield.

**Figure S1:**
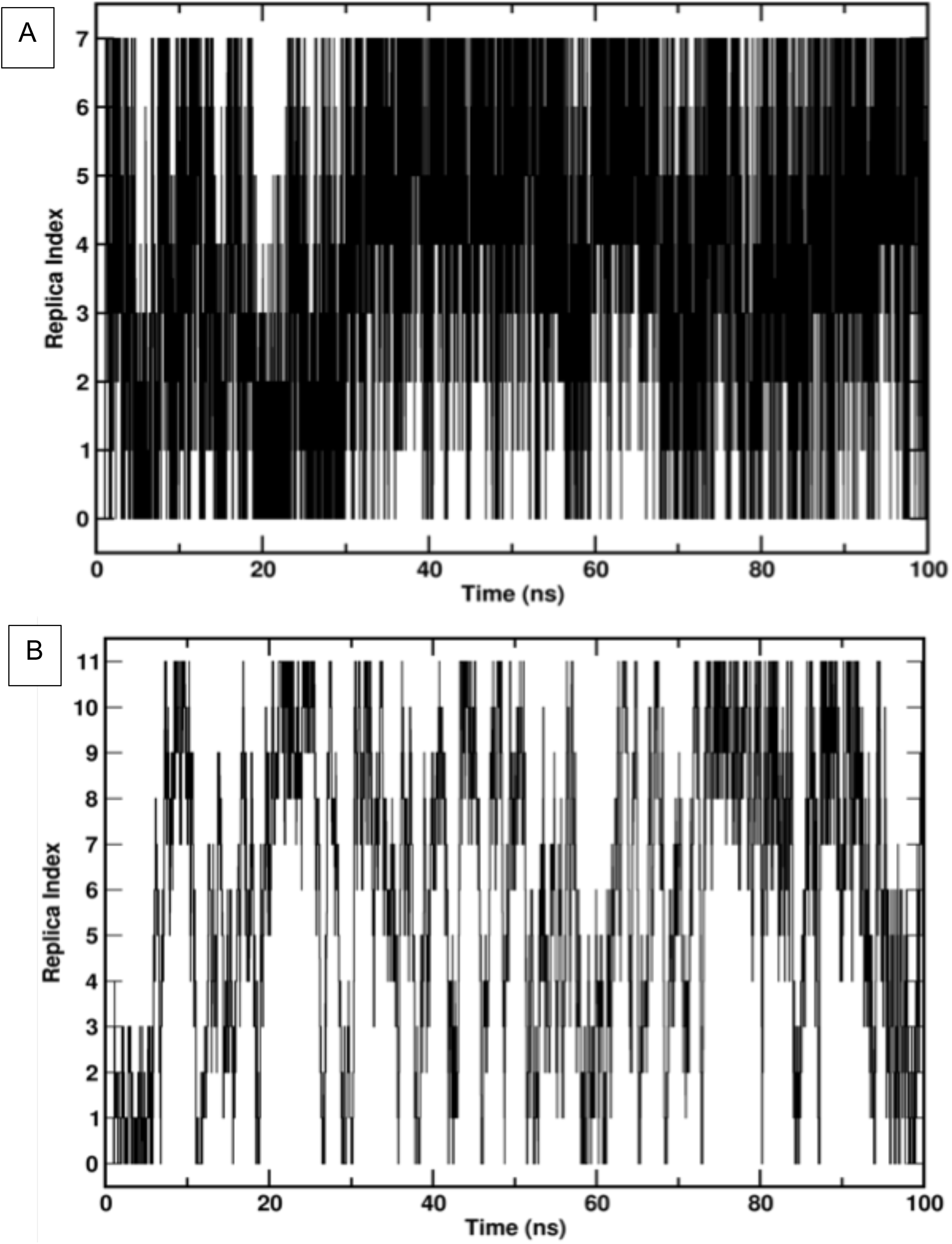
Exchange of base replica (rep 0) across the complete replica state space in TRP-cage system simulated with A) REST2 and B) REHT method. For the sake of clarity only the first 100 ns data are shown.

**Figure S2:**
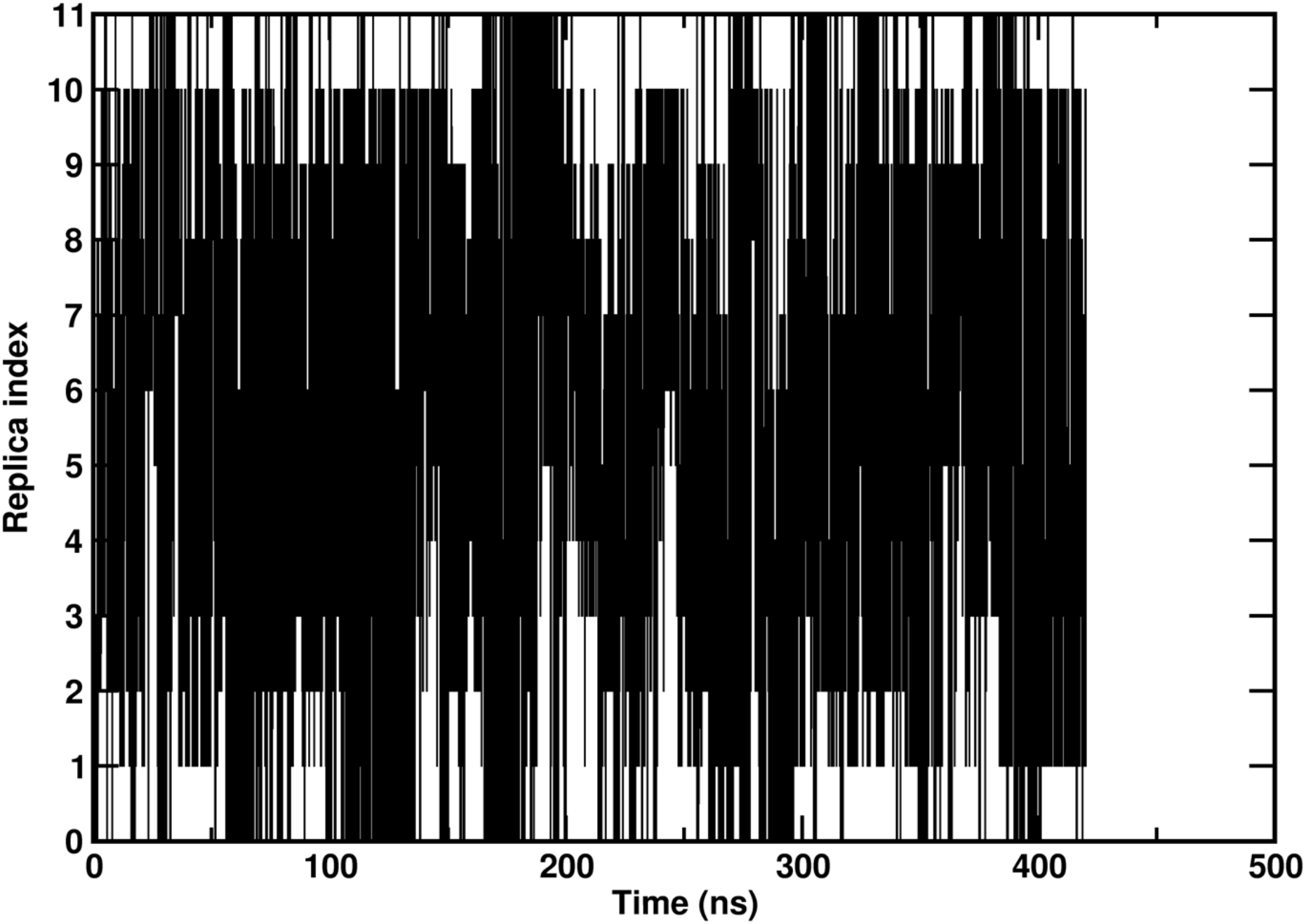
Exchange of base replica (rep 0) across the entire replica state space in beta hairpin system simulated with REHT method. The corresponding simulation of the peptide with the REST2 scheme using the same force field can be found in Ref.^1^

**Figure S3:**
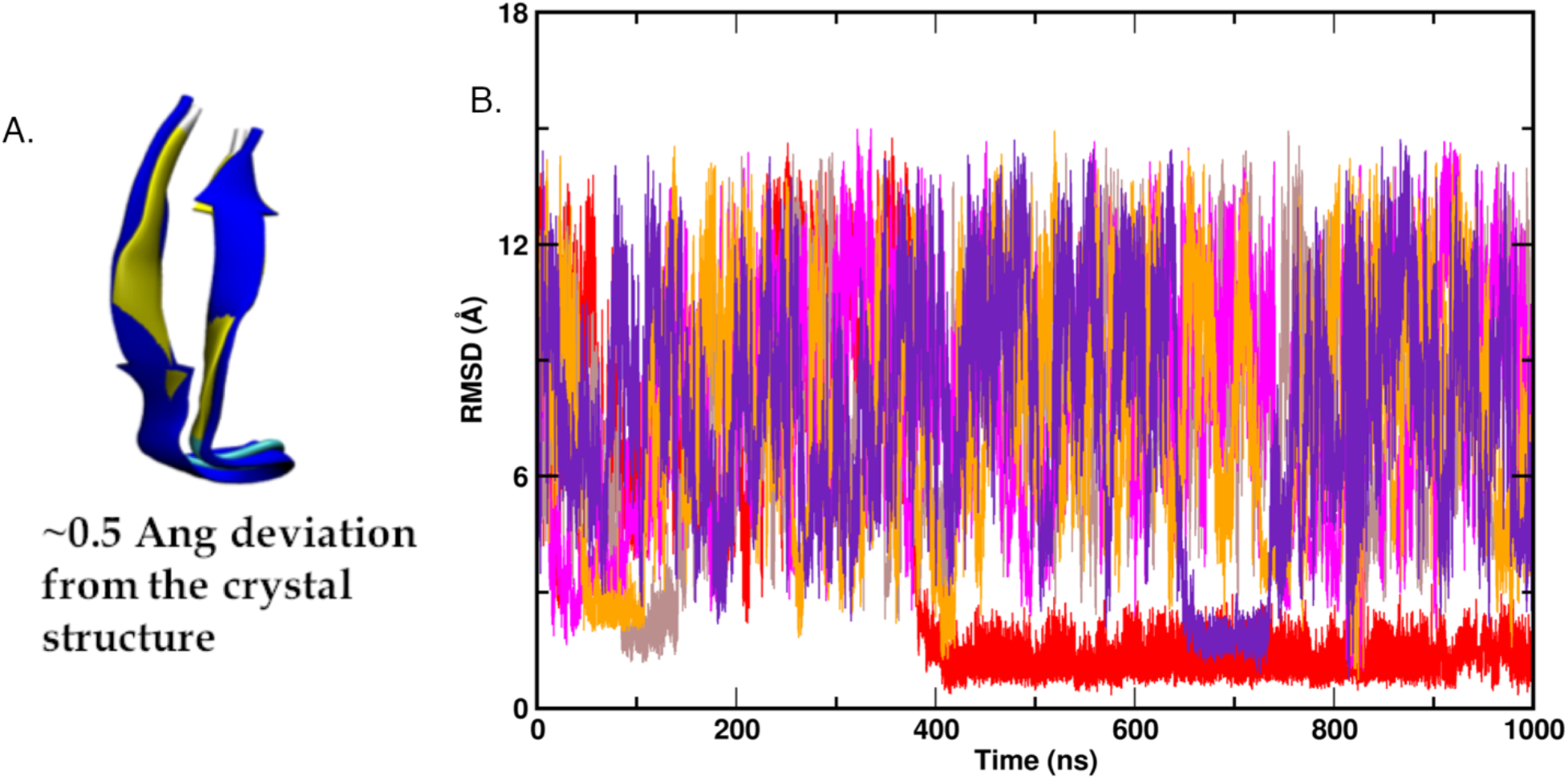
A) Time evolutions of backbone RMSD of REHT-generated Beta hairpin structures with reference to the corresponding NMR structure (1le3), shown for the five replicas that are successfully folded. B) Structural overlay of the simulation-generated beta hairpin (yellow) on the native NMR structure (blue).

**Figure S4:**
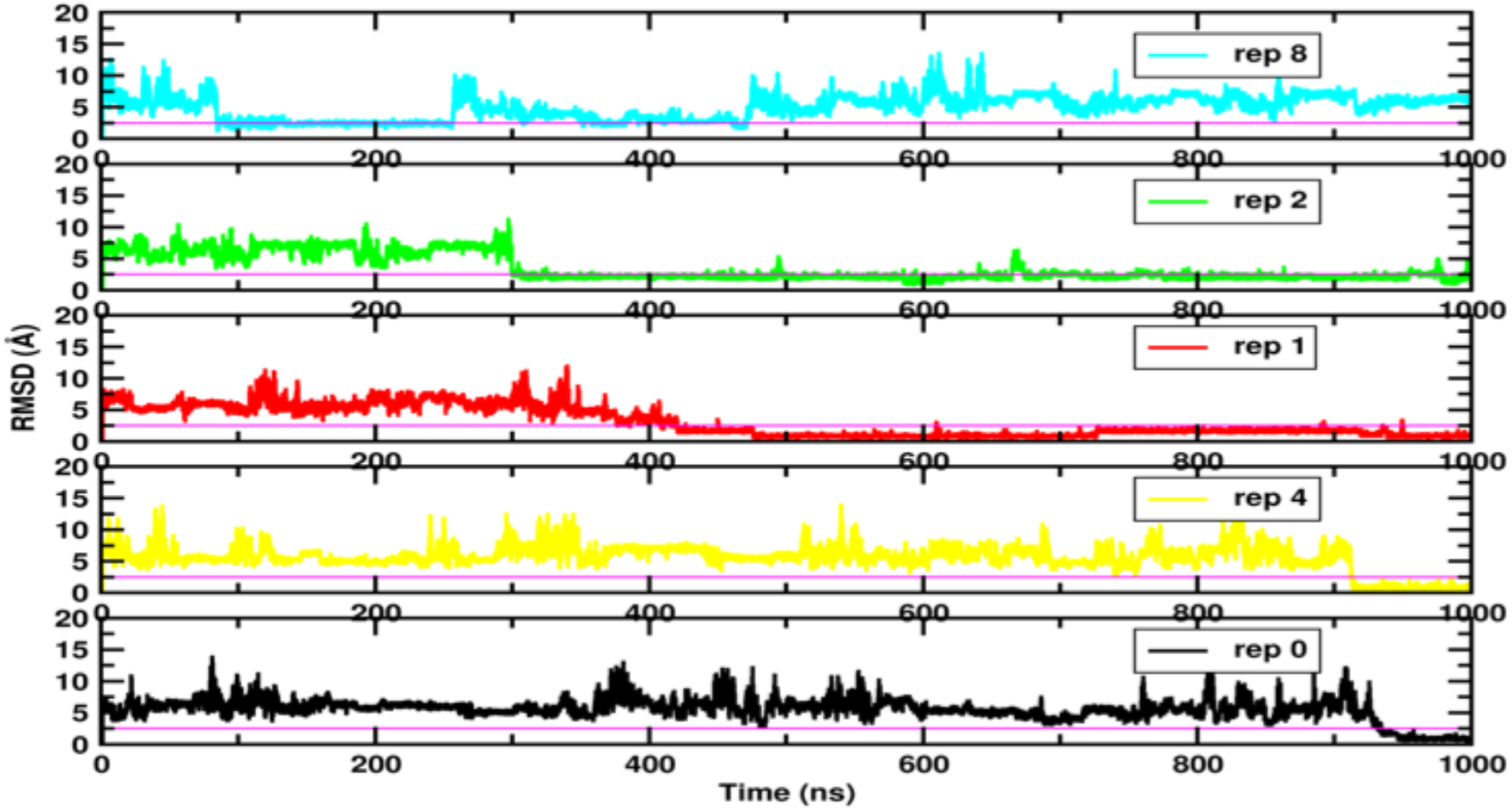
Time evolutions of backbone RMSD of REHT-generated TRP-cage structures with reference to the native NMR structure (1l2y). All the five successfully folded replicas are shown.

**Figure S5:**
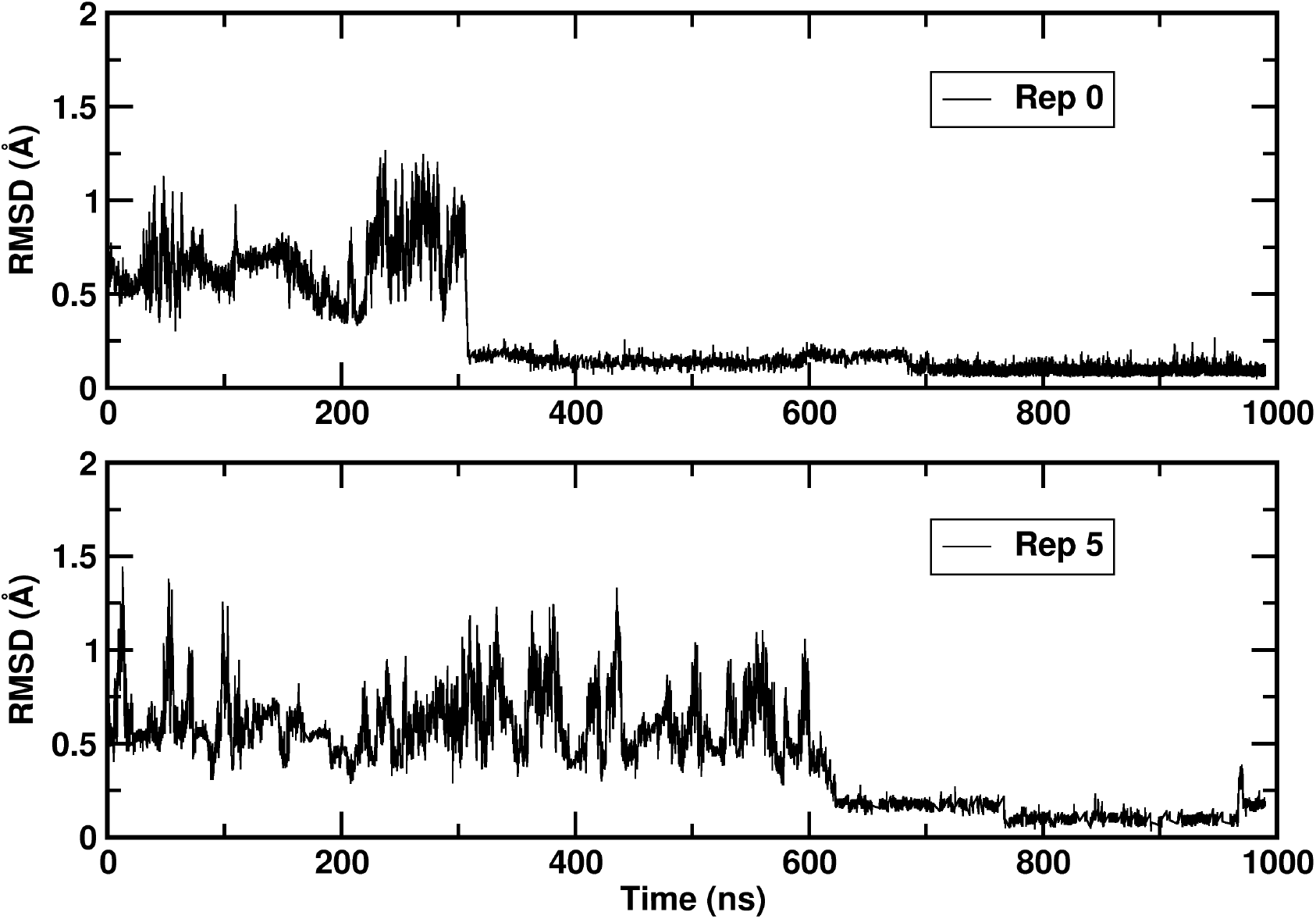
Time evolutions of backbone RMSD of REST2-generated TRP-cage structures with reference to the corresponding NMR structure (1l2y), shown for the only two replicas that are successfully folded.

**Figure S6:**
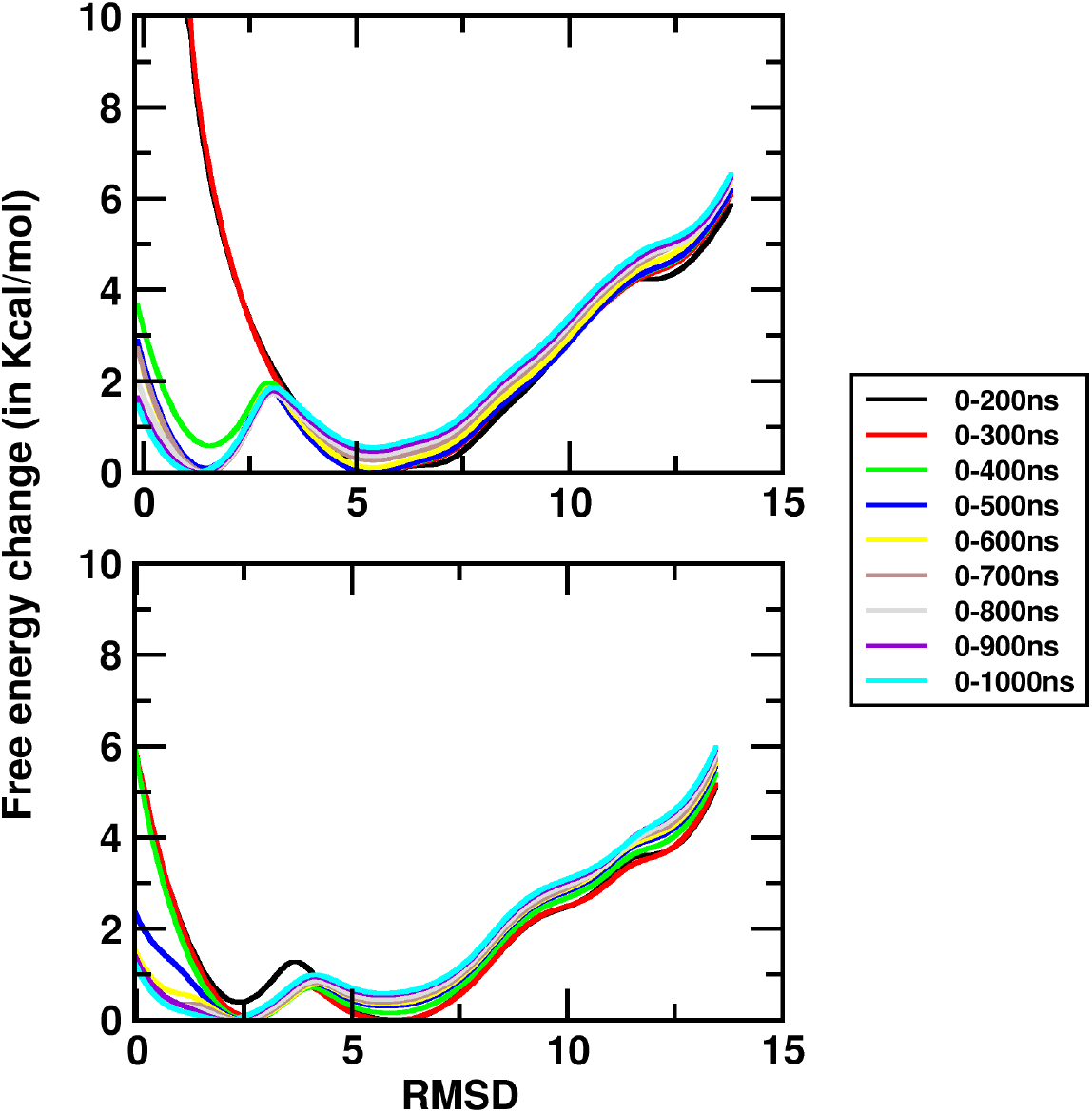
Convergence of free energy landscape of TRP-cage ensemble obtained from A) REST2 and B) REHT simulations. The free energy surface is shown along a single reaction coordinate – RMSD with respect to the native folded NMR structure and plotted as a function of simulation length. The figure clearly indicates that the results obtained from REHT converges faster and better than the REST2 simulations.

## Dihedral switch in Alanine dipeptide

Being the simplest biomolecular model that possesses multiple conformational basins, alanine dipeptide serves as one of the first choices in enhanced sampling literature to study the barrier crossing events. The peptide exhibits three basins across the phi and psi dihedral space. We employed both the conventional REST2 and the novel REHT methods to sample the switching between these dihedral basins of the peptide. The sampled basins across the phi and psi space at different time intervals are illustrated for the lowest temperature replica in Figure S4. While REST2 can sample all the basins within 4 ns timescale, REHT required 6 ns to completely sample this dihedral space. However, the relatively slow transitioning phi angle is frequently sampled in the latter method than the former method (Figure S5). Moreover, the transition was sampled by 4 out of 5 replicas in REHT method, but only once by a single replica out of three in conventional REST2 method. This result suggests that optimal heating of both solute and solvent allows effective crossing of the energy barrier and facilitates frequent transition in the conformational space.

**Figure S7:**
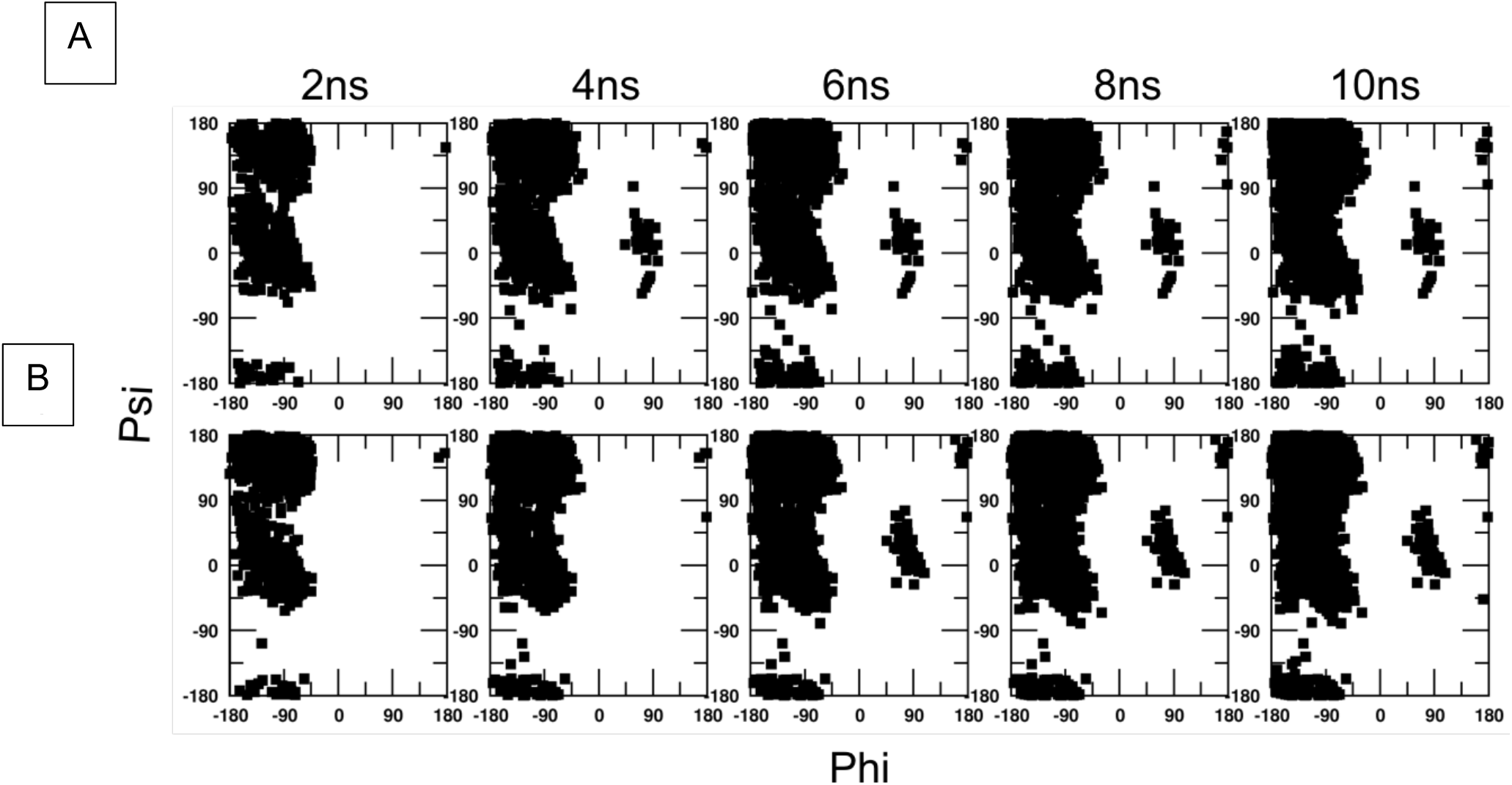
Ramachandran map of alanine dipeptide. The average distribution of dihedral angles of Alanine dipeptide obtained at various time points of REST2 (top panel) and REHT (bottom panel) simulations.

**Figure S8:**
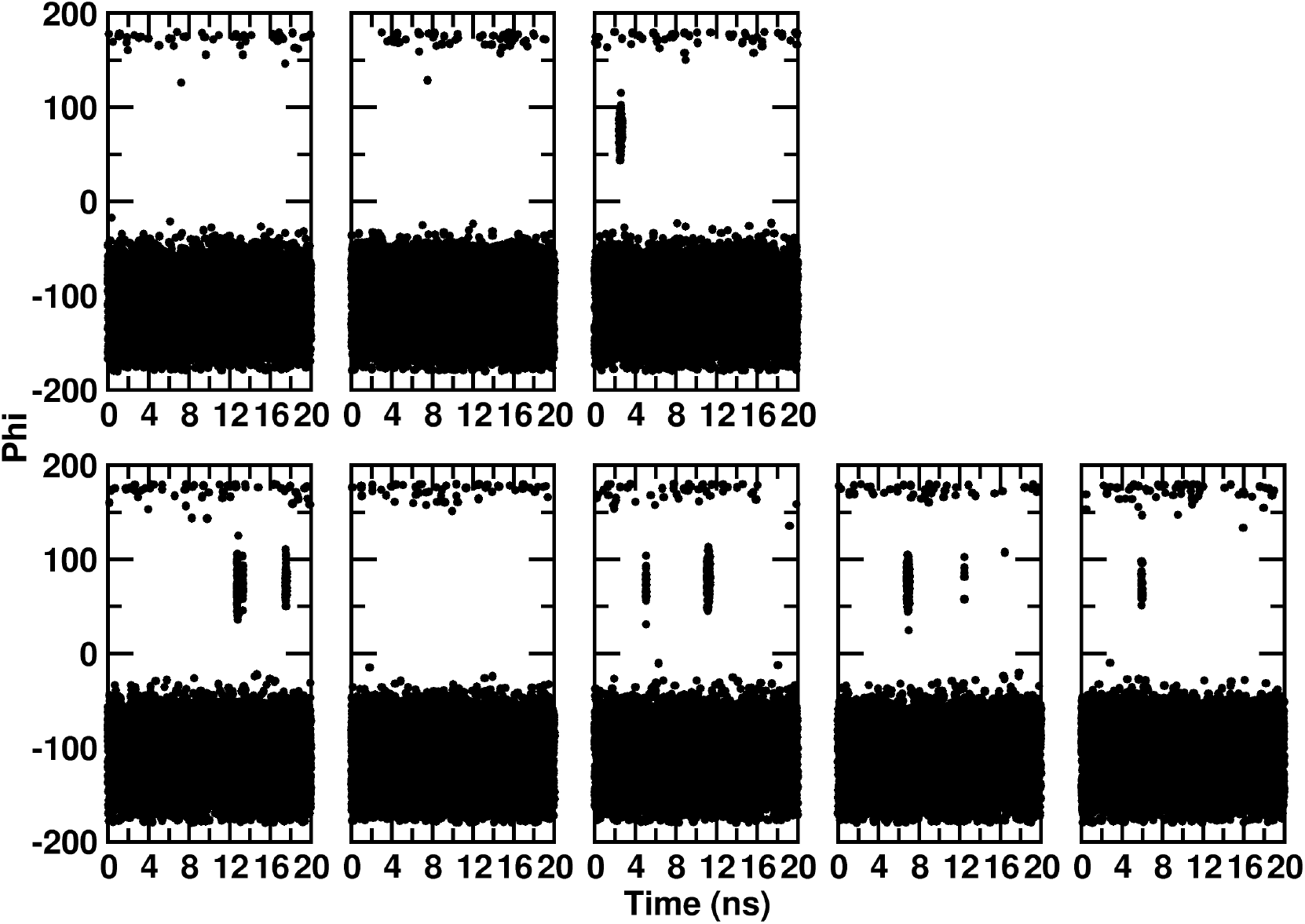
Time evolution of slow transitioning Phi angle of alanine dipeptide simulated using A) three replicas in REST2 simulations and B) five replicas in REHT simulations.

**Figure S9:**
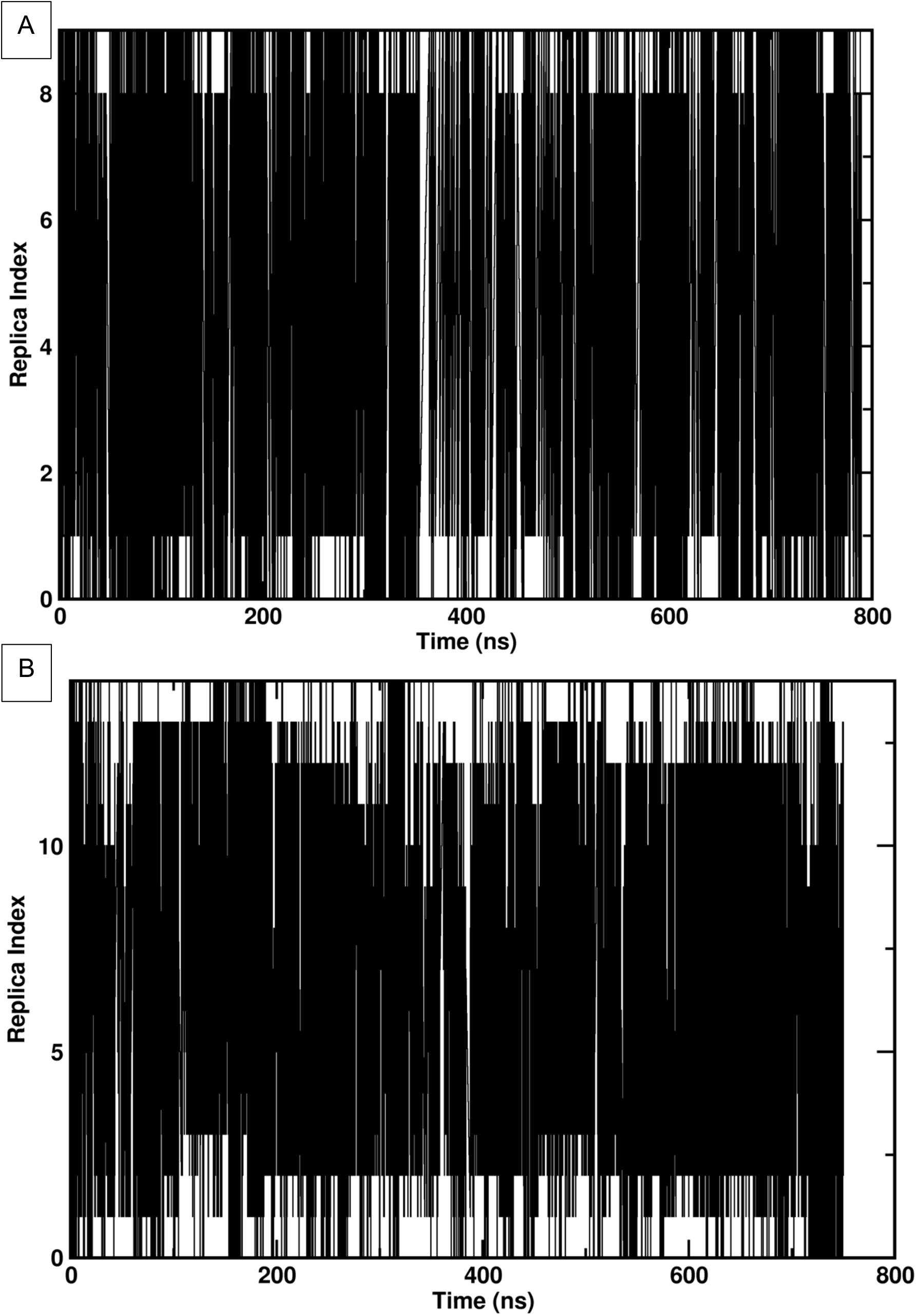
Exchange of base replica (Rep0) of His-5 system across the entire replica space as a function of time in A) REST2 and B) REHT simulation.

**Figure S10:**
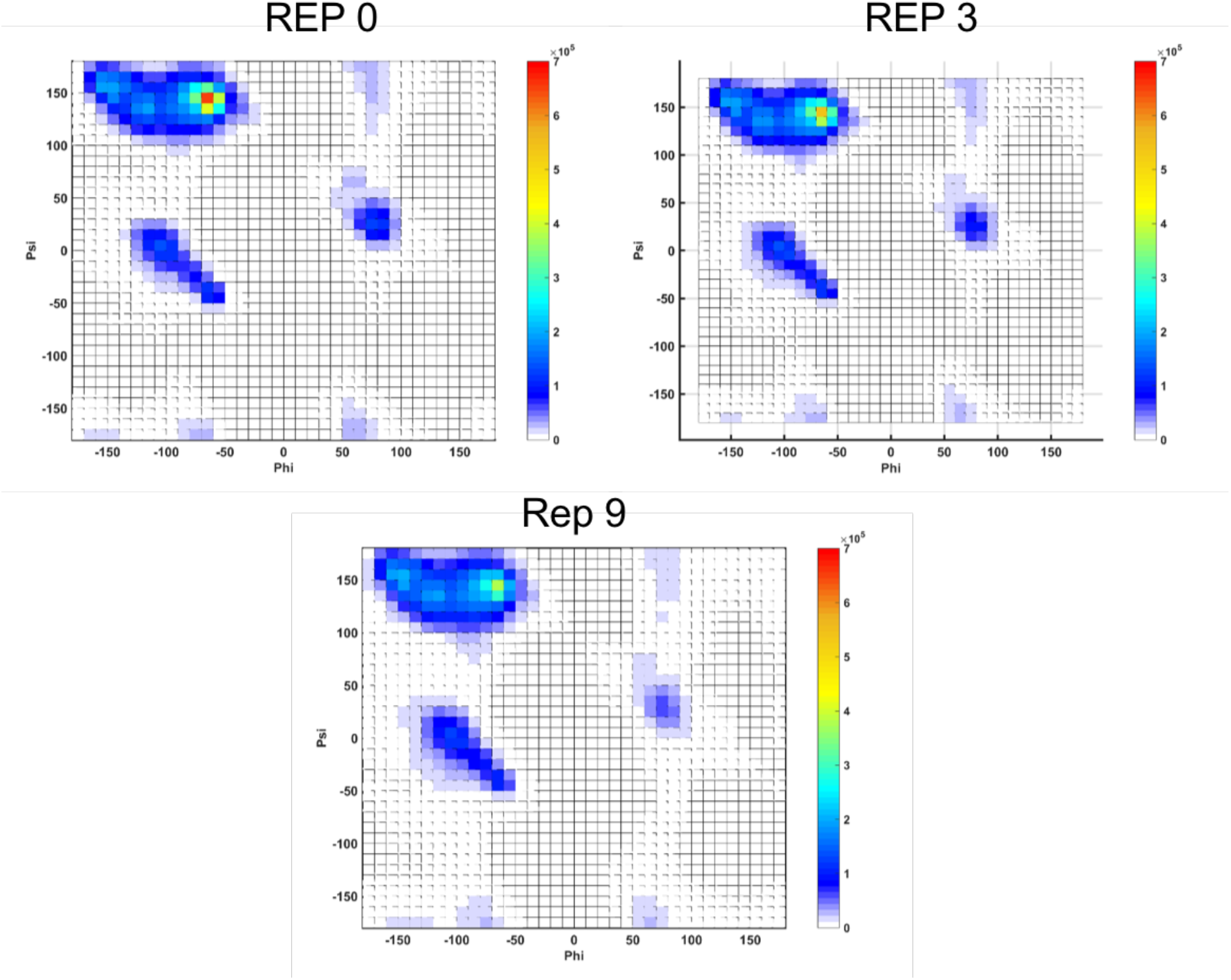
Distribution of dihedral angles projected on the Ramachandran map for all residues of Histatin-5 structures obtaned from REHT simulations. The plot is shown for three replicas (replicas 0, 3 and 9) arranged in increasing order of their respective temperatures (300K, 308K, 325K)

**Figure S11:**
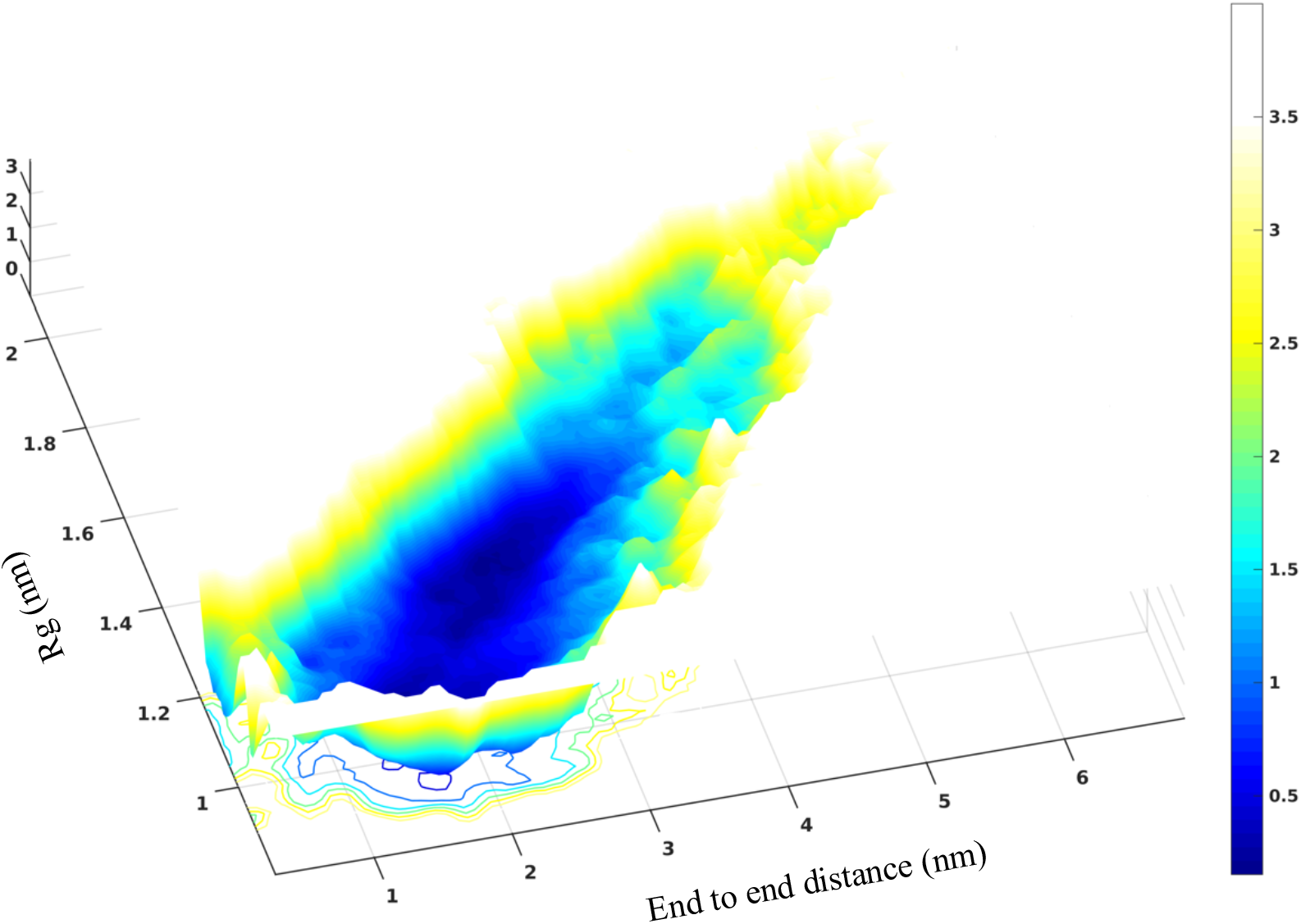
Free energy landscape of Histatin-5 ensemble collected at the base replica of REHT simulation plotted as a function of end to end distance and Rg.

**Figure S12:**
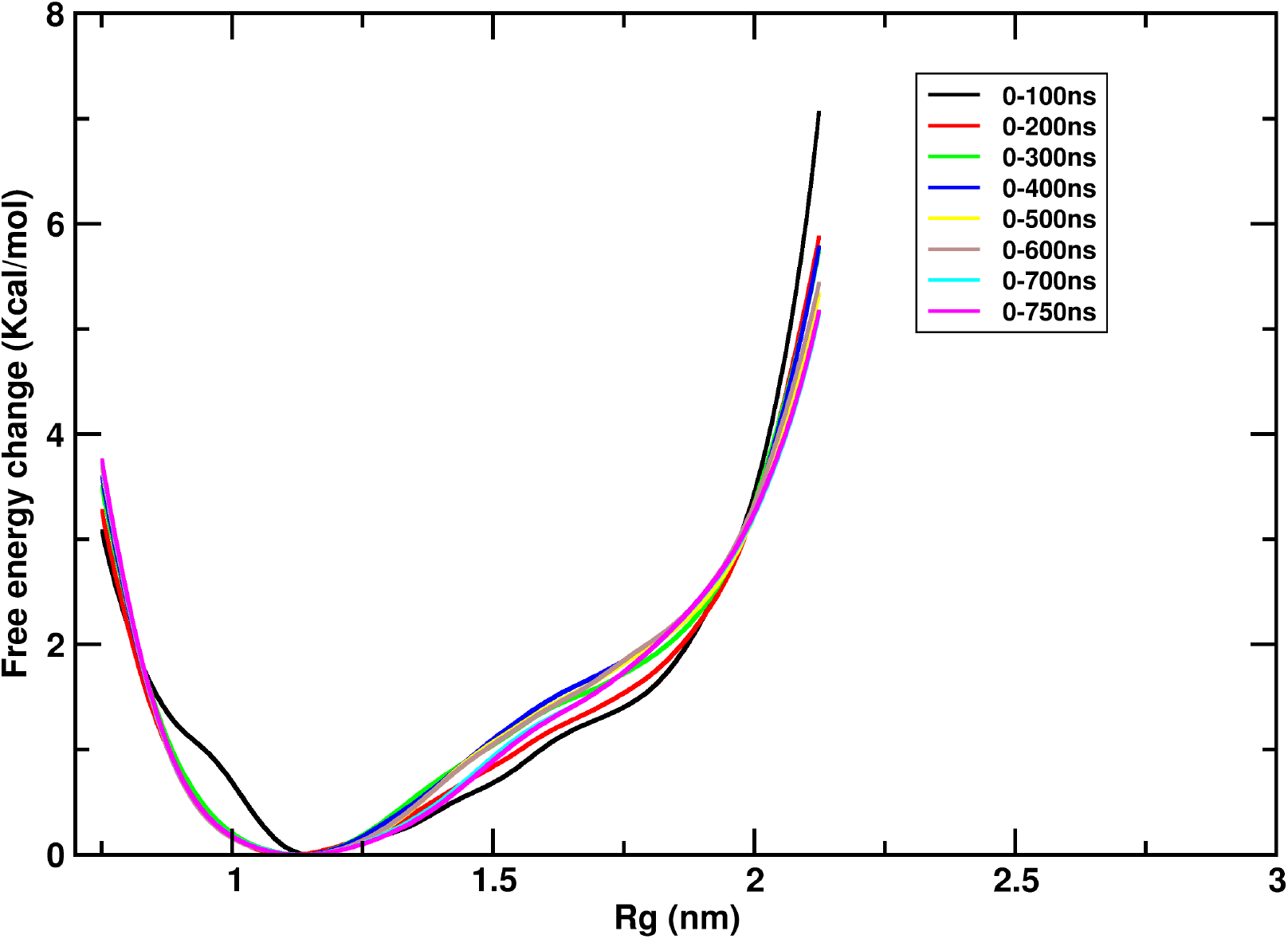
Convergence of His-5 free energy surface along the radius of gyration plotted as a function of simulation length from REHT simulations.

**Figure S13:**
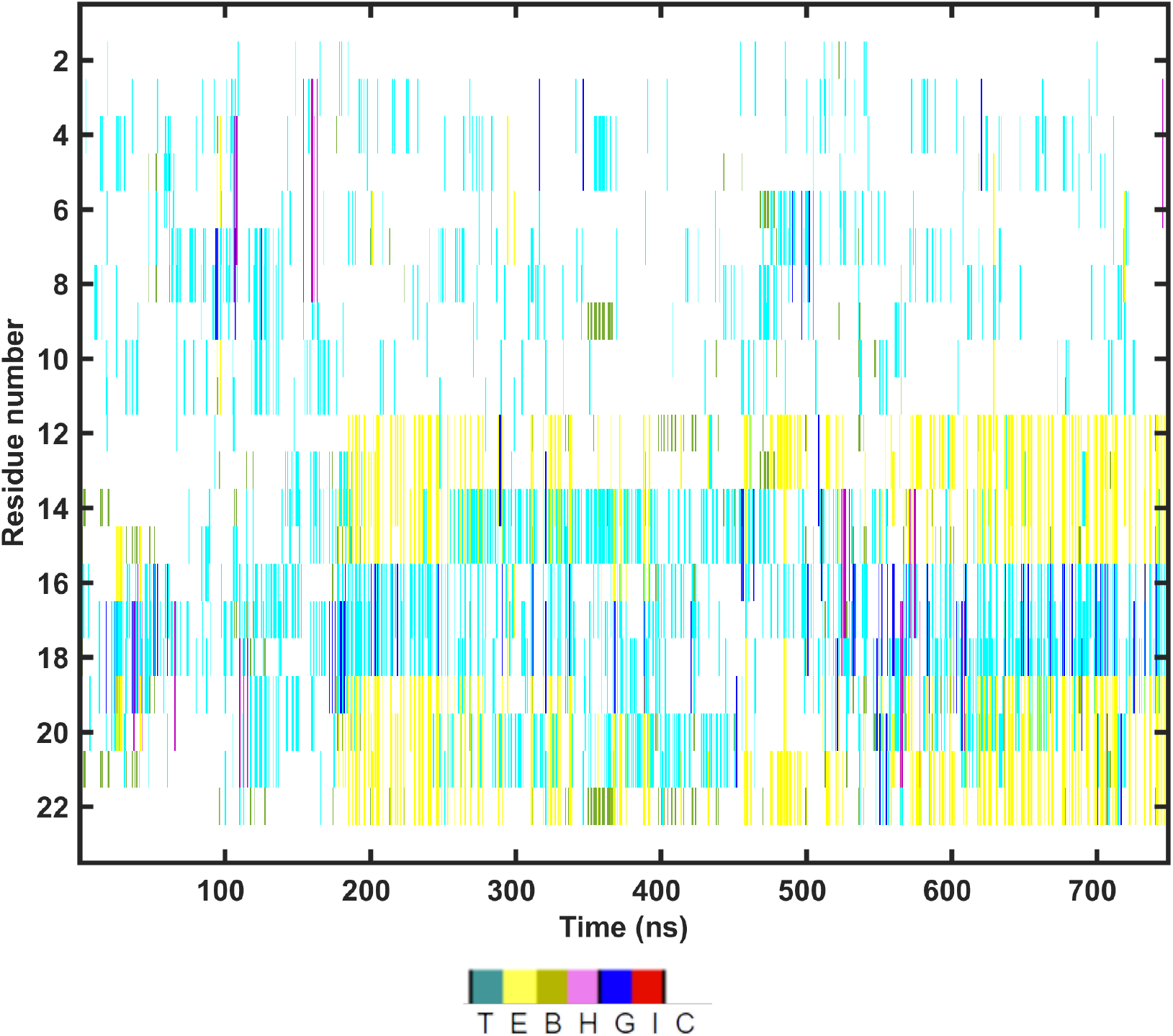
Time series of secondary structure of Histatin-5 ensemble collected at the base replica of REHT simulations.

**Figure S14:**
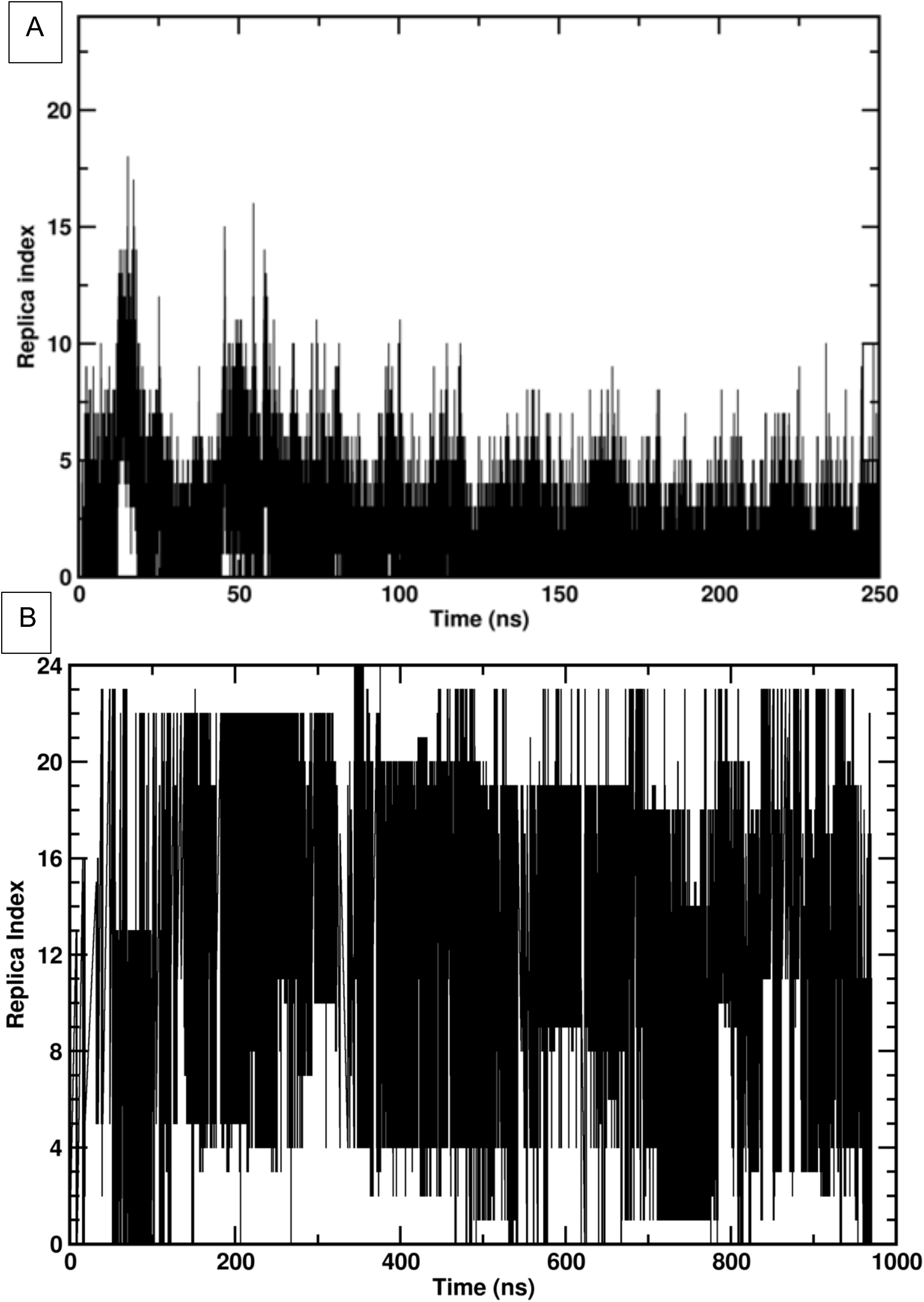
Exchange of lowest rank replica across the complete replica state space in RFA-H system simulated with A) REST2 and B) REHT method.

**Figure S15:**
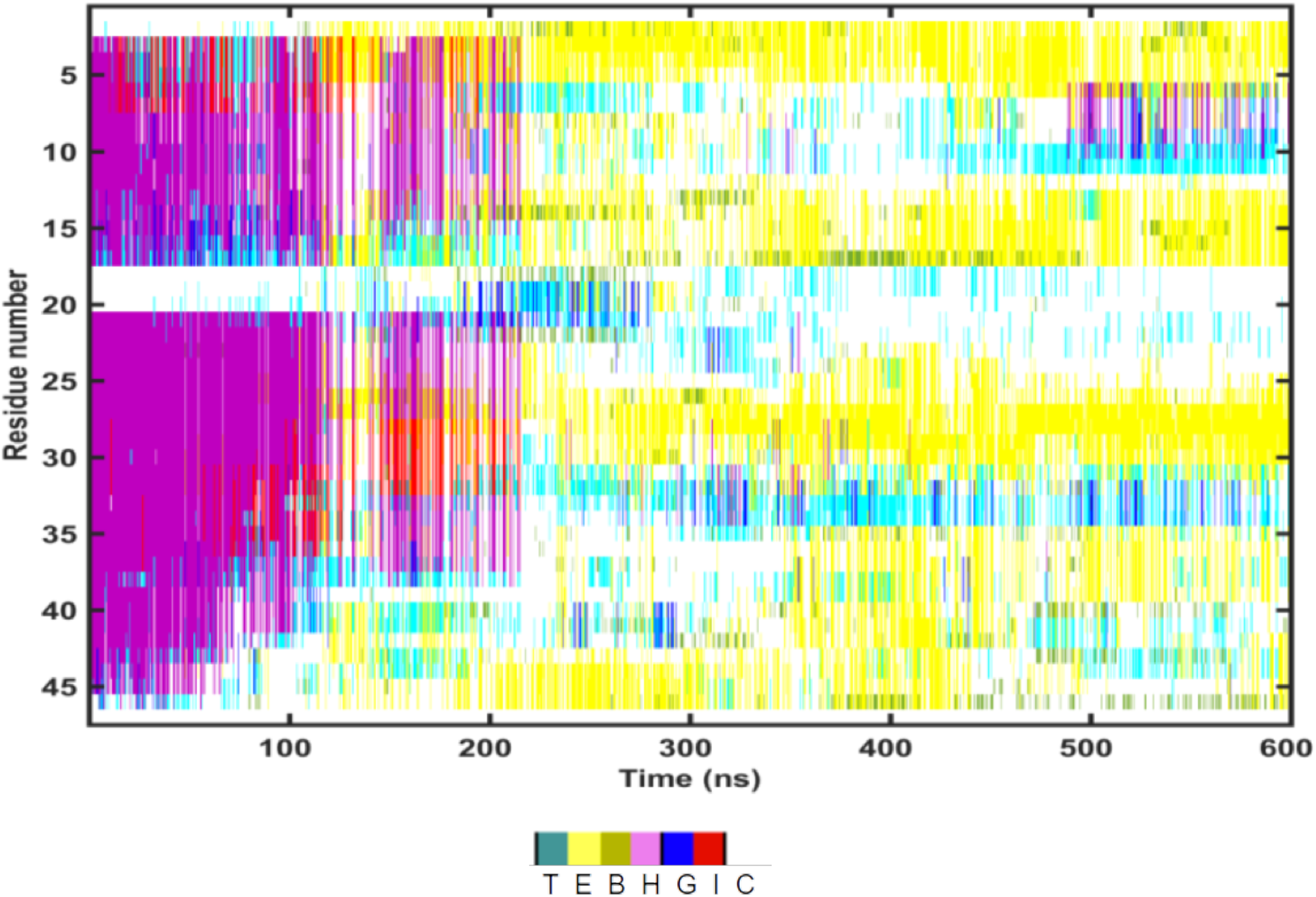
Time evolution of secondary structure for the RFA-H ensemble generated from REHT simulations using all-*α*-helix (2OUG) as starting structure.

**Figure S16:**
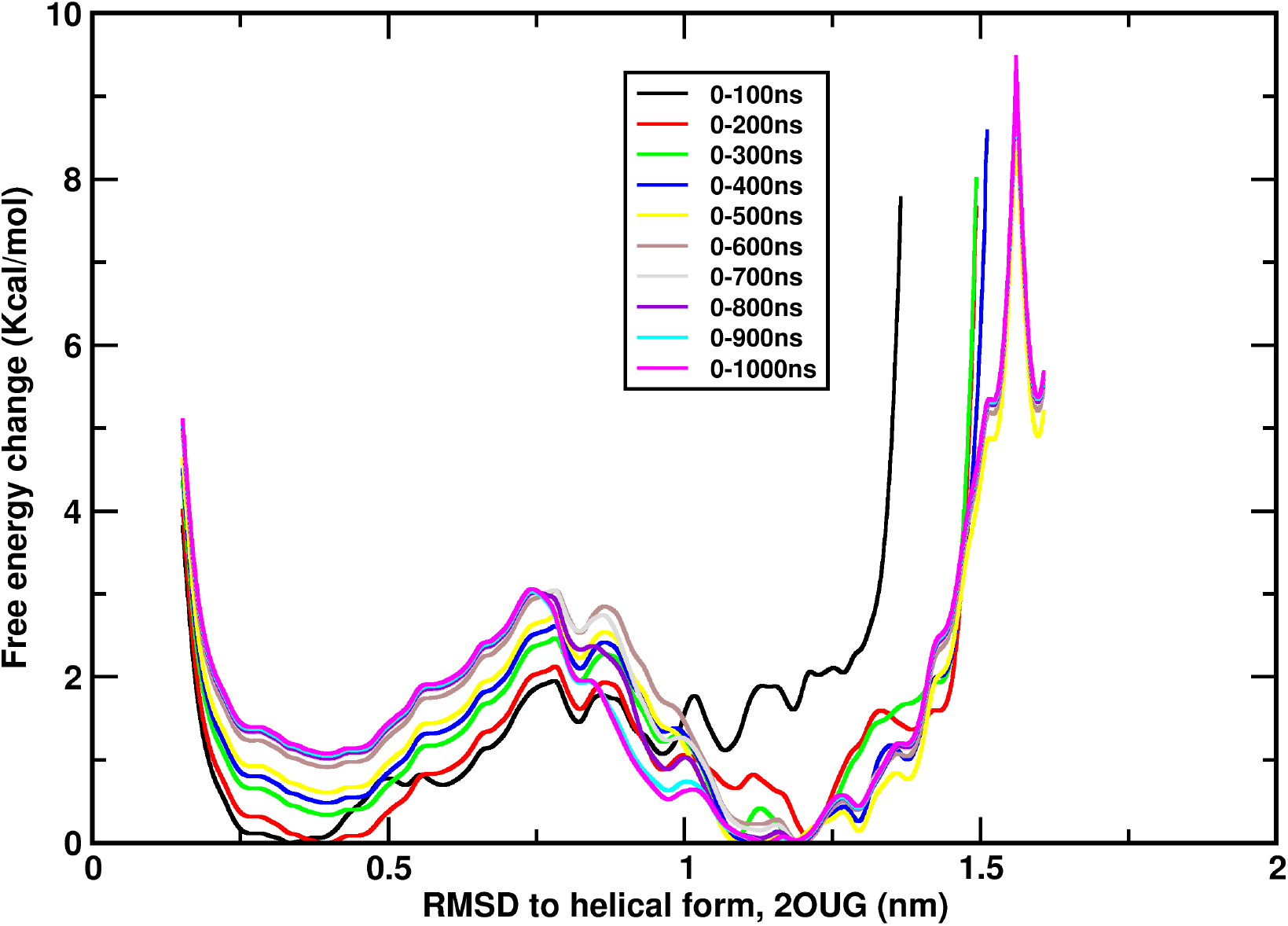
Convergence of free energy surface of RFA-H shown along a single reaction coordinate – RMSD with respect to the helical form (pdb IDL 2OUG). The results are shown for different lengths of REHT simulations.

**Figure S17:**
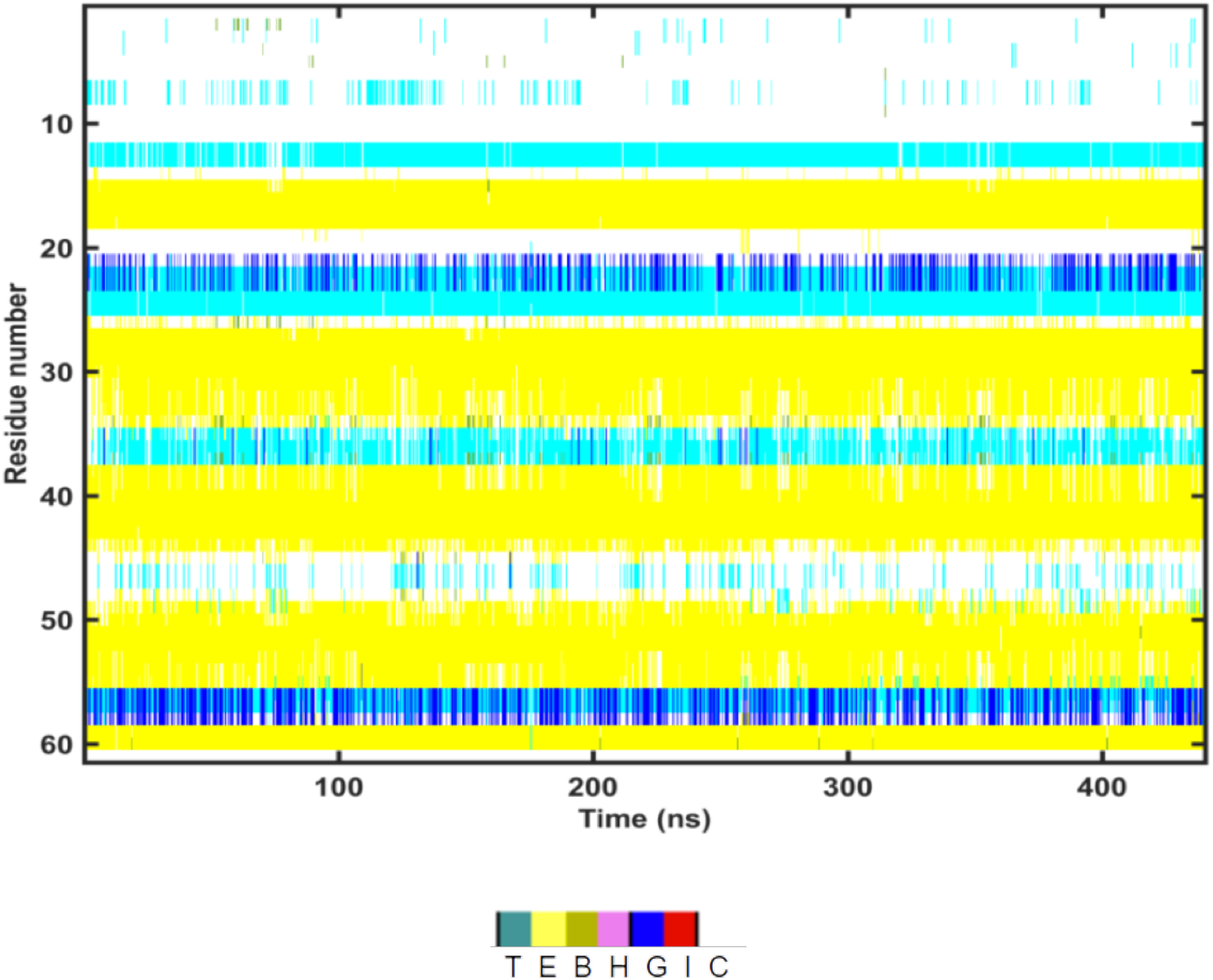
Secondary structure of all residues of RFA-H along the time series. The plot corresponds to the lowest rank ensemble (310K replica) of REHT simulation initiated from the experimental beta barrel structure.

## Replica exchange simulation methodology

Replica exchange simulations allows for studying equilibrium properties with lesser computational time by its rapid relaxation and improved conformational sampling. Originally developed in Monte-Carlo background (Swendsen and Wang, 1986),^2^ the method was later introduced to Molecular dynamics simulations by Sugita and Okomoto in 1999^3^ (Temperature replica exchange molecular dynamics (TREM)). The method achieves effective sampling by simulating a series of low and high temperature replicas, while allowing the exchange of configurations at regular intervals. Moreover, due to its stochastic nature of exchanges that ensures the detailed balance, it generates Boltzmann weighted ensemble from which it is straight forward to obtain the thermodynamic averages. The probability of accepting the exchange between replica m and n depends on the difference in the Boltzmann weight factor, that is exponentially related to the difference in energy and temperature (Eqn.1).

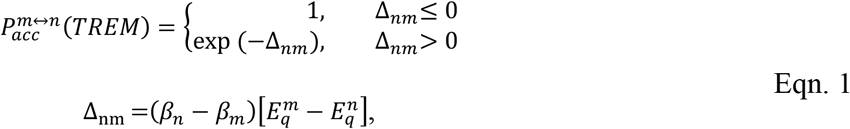

Where *β*_*n*_ and *β*_*m*_ are reciprocal temperatures (1/(*K*_*B*_*T*_*n*_) and 1/(*K*_*B*_*T*_*m*_) respectively). 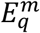 and 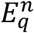 are potential energy of replicas m and n respectively. For a larger system the temperature differences between two replicas should be chosen minimal to yield viable exchange acceptance ratio. This demands large number of replicas to cover a sufficient range of temperatures.

### 3.1.1 Replica exchange solute tempering (REST2)

Instead of changing the temperature across the replica ladder the advanced replica exchange solute tempering (REST2) method scales the energy function in a particle-wise manner such that the solute is effectively heated up while keeping the water cold. For instance, the potential energy function of replica m is broken down into intramolecular protein interactions (*H*_*pp*_), protein-water interactions (*H*_*pw*_), and water self-interactions (*H*_*ww*_) whose potentials are scaled as shown in Eqn. 2.

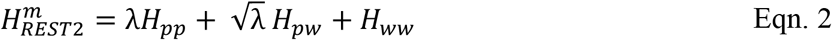

Where 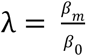, in which *β*_0_ and *β*_*m*_ are reciprocal temperatures of 0^th^ and m^th^ replicas. While running all the replicas at the same temperature, the method cancels out the energy difference in water self-interaction energy that otherwise hugely contribute for the poor scaling as in TREM. Thus, the acceptance probability of REST2 as shown in Eqn. 3 depends only on the energy differences of intramolecular solute energy and intermolecular energy between protein and water.

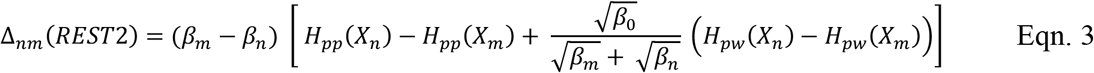

Where, *H*_*pp*_(*X*_*m*_) and *H*_*pp*_(*X*_*n*_) indicates the intramolecular energy of protein solute in m^th^ and n^th^ replicas. *H*_*pw*_(*X*_*m*_) and *H*_*pw*_(*X*_*n*_) are interaction energies of protein and water in the two replicas. *β*_*m*_, *β*_*n*_ and *β*_0_ are reciprocal temperatures of replicas m, n and 0^th^ replicas respectively.

### 3.1.2 Replica exchange hybrid tempering (REHT)

Though the REST2 claims to be efficient for studying larger proteins including that of IDPs as shown recently by Shrestha et.al, it suffers from inability to explore the complex energy landscape with larger energy barriers. We speculated that this could be due to the imbalance between hot solute and cold solvent that causes differential dynamics of central protein and surrounding bulk water. In general, the biomolecular folding and conformational transition is tightly coupled to its surrounding water dynamics. Hence in this work, we introduce a hybrid method that optimally treats the protein as well as the surrounding water. We achieve this by associating the replicas to different bath temperatures in addition to scaling down the potential function of protein in contrary to REST2. Treating the replicas in such a combination doesn’t violate the detailed balance condition. More importantly the method yields expedited protein conformational sampling by allowing efficient rewiring of water. For the viable exchanges the temperature gaps between the adjacent replicas are kept minimal. At the same time the protein is allowed to effectively heated-up to a larger extent by additional REST2 scaling factor.

## Derivation of REHT method

Let us consider the collection of replicas simulated in replica exchange simulation as {X1, X2…Xn}. In general, the replicas differ in temperature (Ti) while uses an identical Hamiltonian function (Temperature replica exchange) or vice versa (Hamiltonian replica exchange). In our hybrid approach, we change both the temperature as well as the Hamiltonian across the replicas. Hence the replicas can be denoted as {*X*_*m*_, *H*_*m*_(*X*_*m*_), *T*_*m*_}, where *X*_*m*_, *H*_*m*_(*X*_*m*_), *and T*_*m*_ respectively are configuration, potential energy function and temperature of replica m.

Since the replicas are non-interacting, the equilibrium probability of this larger ensemble can simply be obtained by the product of Boltzmann factors of each replica.

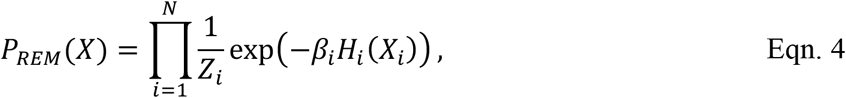

Where, *β*_*i*_ denotes the inverse temperature (1/*K*_*B*_*T*) and *Z*_*i*_ represents the configurational partition function.

Consider an exchange of configurations between a pair of replicas m and n.

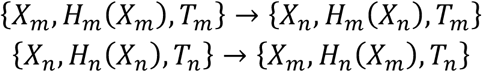

The corresponding transition probability for this exchange (*X*_*m*_ → *X*_*n*_) would be given by,

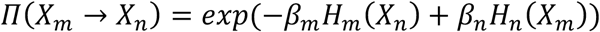

Imposing the detailed balance condition, where the reverse exchange is allowed with equal probability,

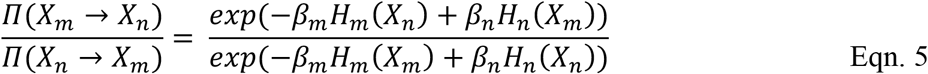

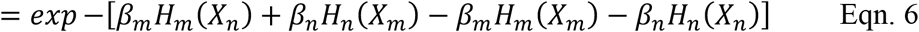

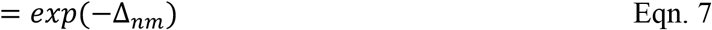

Where Δ_*nm*_

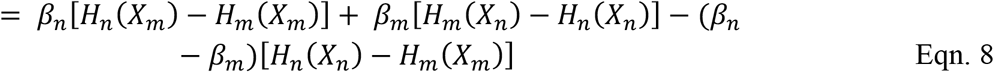

With the Metropolis criteria, the probability of accepting the exchange X_m_ → *X*_*n*_ becomes,

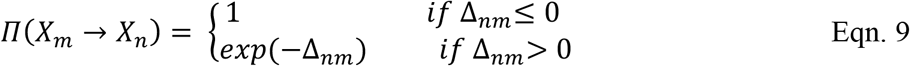

We used a deformed Hamiltonian identical to the one used in the REST2 approach.

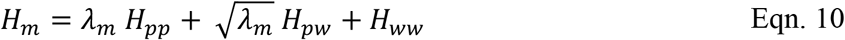

Applying Eqn.10 on Eqn.8, Δ_*nm*_ becomes

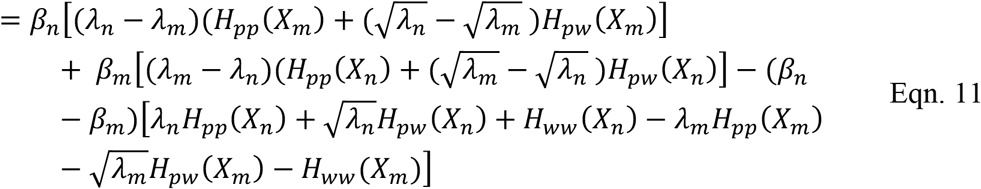

Coefficient of *H*_*pp*_:

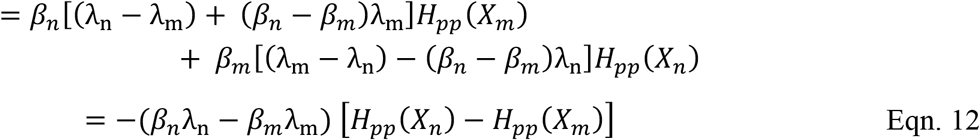

Coefficient of *H*_*pw*_:

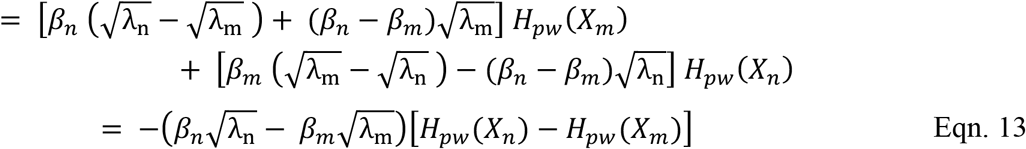

Coefficient of *H*_*ww*_:

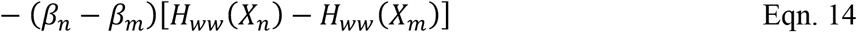

Combining Eqn. 12, 13 and 14,

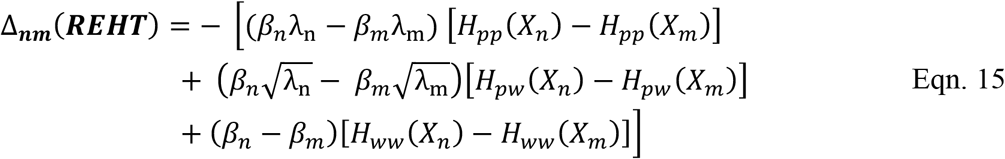

## Notes

### Competing Interest Statement

The authors have declared no competing interest.

